# Rapid ecological and evolutionary divergence during a poleward range expansion

**DOI:** 10.1101/2021.11.04.467261

**Authors:** Monica M. Sheffer, Brian Schulze, Linda Zander, Pierick Mouginot, Thomas Naef, Michael Lalk, Martina Wurster, Clara Ahrend, Jürgen Kreyling, Rosemary G. Gillespie, Katharina J. Hoff, Stefan Prost, Henrik Krehenwinkel, Gabriele Uhl

## Abstract

In response to climate change, a northward range expansion has been observed in many species. The wasp spider, *Argiope bruennichi*, has expanded from its historic range in the Mediterranean (“core”), now reaching as far as the Baltic States and Scandinavia (“edge”), even faster than the pace of climate change. We explored life history traits, adult phenotypes, offspring cold tolerance, and genomic patterns across the European range of *A. bruennichi*, and found origin-, environment- and life stage-specific responses to the cold northern climate. Wasp spiders have shifted their phenology at the edge, with females maturing earlier and at a smaller size, but maintaining similar pigmentation, clutch sizes, and hatching success compared to the core region. Using a reciprocal common garden experiment on overwintering offspring from the core and edge, we found evidence for genetic adaptation and considerable phenotypic plasticity. Overwintering survival was lower under the cold winter treatment for spiderlings from both origins. However, the edge-origin spiderlings that survived the winter had lower lethal temperatures and enhanced supercooling ability with reduced phenotypic plasticity in supercooling points compared to core spiderlings, while the chill coma recovery time was similar. Metabolomic analysis revealed accumulations of amino acids and myo-inositol in the cold winter treatment, particularly in spiderlings from the edge population, suggesting a role of these metabolites in improving cold tolerance. Genotype-environment tests showed strong genetic association across the genome to seasonality and minimum winter temperature. The population genomic analysis across the European range splits *A. bruennichi* into two distinct genetic clusters through the center of Germany, which roughly aligns with turnover from an oceanic to continental climate zone, complementing the genotype-environment test results. Overall, our study highlights the importance of integrating data on phenological shifts, changes in life-history, and life stage-specific phenotypic plasticity and genetic adaptation to understand the impacts of range expansions and shifts. The nuanced processes of acclimation and adaptation we uncovered advocate for holistic investigations of evolutionary fitness and fitness-related traits in the context of organismal responses to novel and changing environments.

## Introduction

Global climate change is a major stressor for ecosystems, leading to extinction events, dwindling population sizes, range contractions, or range shifts in many species worldwide (Harvey et al., 2023; Pecl et al., 2017; Wagner, Fox, et al., 2021; Wagner, Grames, et al., 2021; Warren et al., 2018). In the face of changing environments, some organisms can disperse in search of suitable habitats or show *in situ* responses via phenotypic plasticity within a single generation and/or via genetic adaptation over successive generations. Given the observed rate of climate change, which often outpaces the dispersal capacity of many organisms (reviewed in Lenoir et al., 2024), we expect that species that show a combination of movement, phenotypic plasticity, and genetic adaptation will likely be the species that have the potential to persist.

A habitat beyond the original range of a species might become suitable in one dimension, such as providing sufficiently warmer summer temperatures for development, while other dimensions, such as colder winter temperatures, higher humidity or differences in diel light cycles, may strongly limit expansion and range shifts (Ljungström et al., 2021). In this novel environment, local genetic adaptation may arise if selection pressures are high and there is sufficient genetic variation. This has, for example, been well documented in range-expanding damselflies, which show local adaptation in cold tolerance and genetic differentiation at the northern expanding edge (Carbonell et al., 2021; Dudaniec et al., 2018; Lancaster et al., 2015, 2016).

To understand the processes that allow for establishment in novel environments, we are studying a range-expanding spider species, *Argiope bruennichi,* which has undergone a well-documented latitudinal range expansion within Europe in less than a century. While its European range was mostly restricted to the Mediterranean before 1960, it rapidly expanded as far as Germany, Poland, and the southern UK in the 1990s (Barabasz-Krasny & Andrzej, 1998; Guttmann, 1979; Smithers, 2000; Wawer et al., 2017). Continuing northward from then on, it arrived in coastal parts of Scandinavia in late 1980s and early 1990s (Follner & Klarenberg, 1995; Jonsson, 2004; Jonsson & Wilander, 1999; Scharff & Langemark, 1997), establishing there in the early 2000s (Bratli & Hansen, 2004) and finally becoming established as far north as Estonia in 2006 (Algo, 2010; Krehenwinkel & Tautz, 2013; Kumschick et al., 2011). This rapid range expansion has outpaced climate change, in that the species now inhabits areas that are much colder than its original range (Krehenwinkel et al., 2015). Given this remarkable range expansion, we use *A. bruennichi* as a model system to study two overarching questions: 1) How does this species cope with the shorter growing season and the much colder winter temperatures of northeastern Europe – through phenotypic plasticity, local adaptation, or a combination? And if we uncover evidence suggesting local adaptation, and if populations are genetically differentiated, 2) does genetic differentiation correlate with environmental variation, corroborating a conclusion of local adaptation? Understanding the processes that allowed this species to rapidly establish in northern latitudes will provide insight into how organisms can respond as global temperatures change and range shifts become more common.

Tests of the genetic basis of phenotypic traits can be performed using common garden or reciprocal transplant experiments, which allow phenotypic variation to be partitioned into the relative contributions of genetic and environmental sources. In an ideal system, multiple generations could be reared in the laboratory to ensure that transgenerational plastic or “maternal” effects are not conflated with genetic adaptation. However, many species that show interesting distributional changes cannot be reared in the laboratory, as is the case with *A. bruennichi*. Therefore, we combine phenotypic evidence from natural populations and experimental approaches with genomic evidence to offer insight into the degree of phenotypic plasticity and possible genetic adaptation in this system. These approaches are rarely used in concert, especially across different life stages, as is done here.

We expect *A. bruennichi* spiders from northeastern (“edge”) populations to have traits that enable them to survive and reproduce despite colder winter temperatures, cooler summers, and a shorter growing season relative to the ancestral “core” of their range. Given the pace of the range expansion, we expect that these spiders display considerable phenotypic plasticity in these traits, which may have enabled initial colonization. However, as the range expansion has progressed over the course of several decades and many generations, it is possible that we may detect genetic adaptation that has alleviated fitness constraints imposed by the novel environment. We investigated a suite of life history and cold tolerance traits that we expect may be under selection and play a role in the successful colonization and establishment of the wasp spider in northeastern Europe. We focused on two phases of the life-cycle of *A. bruennichi* (**Supplementary Figure S1**): the reproductive season in the summer and the overwintering season of the juveniles (“spiderlings”) in the egg sacs. Considering these two life stages allows us to investigate traits relevant to different components of fitness: fecundity and survival.

The traits investigated for the reproductive season relate to adult females that were collected across a transect of their range from France to Estonia. We explored in what way adult females along the transect differ in phenology, body size, fecundity, and reproductive output. Due to the later spring and cooler temperatures in the north, we expected spiders from the edge of the distribution to develop more slowly and therefore mature later compared to those from the core area. Alternatively, spiders might mature early, but at a smaller size due to the restricted season length and in accordance with the converse Bergmann’s rule (Blanckenhorn & Demont, 2004; Masaki, 1967). Shorter summers with increasing latitude may further limit reproductive success by constraining accumulation of resources for egg production, leading to later oviposition, smaller clutch sizes with possibly smaller spiderlings (but see Wolz et al., 2020) and possibly reduced hatching success. We also investigated if females are darker pigmented with increasing latitude, due to a positive thermoregulatory effect of melanism in an environment with shorter summers and overall lower temperatures (Yavad et al., 2018).

To determine how *A. bruennichi* copes with environmental challenges during the overwintering season and to assess the degree of plasticity and genetic adaptation in survival-related traits, we performed a reciprocal common garden experiment, investigating differential overwintering survival, body size, mass and several cold tolerance traits in juvenile *A. bruennichi*. Arthropods display several traits that allow them to survive harsh winter temperatures, dependent upon their cold tolerance strategy. The cold tolerance strategy of only a few spiders has been assessed, and all were classified as freeze avoidant, e.g. they die upon internal ice formation (Anthony et al., 2019; Cubillos et al., 2018; Kirchner, 1973, 1987). Under the assumption of genetic adaptation to colder climate at the edge of the range, we formulated several predictions. We predicted that spiderlings from the edge of the range would be smaller as we hypothesized that smaller size and mass would be advantageous as it would reduce the probability of spontaneous ice formation. We further predicted that spiderlings from the edge have lower supercooling points, lower lethal temperatures and shorter chill coma recovery times. We explored the underlying metabolomic changes and predicted that spiderlings from the edge would be more likely to accumulate cryoprotectants due to harsher winters and accumulate metabolic intermediates due to reduced metabolic rates. However, under the assumption of phenotypic plasticity, spiderlings from the core and edge of the range would show similar response patterns.

We investigated the degree of genetic differentiation in adult *A. bruennichi*, drawing on previous population genetic studies which suggested that rapid range-expansion was driven by adaptive introgression of previously isolated eastern lineages, providing alleles preadapted to colder climates and enabling rapid adaptation to northern climates (Krehenwinkel et al., 2015, 2016; Krehenwinkel & Tautz, 2013). In the current study, we focus on the European range of the species, covering populations from the Mediterranean core of the distribution to the expanding northern populations in the Baltic states. We expected a turnover from “ancestral” to “expanding” genotypes in Germany and predicted a broad genetic cline, as the spider is able to disperse by ballooning with a silken thread (Follner & Klarenberg, 1995; Wolz et al., 2020). Further, we would expect to see strong correlations of genotype frequencies with environmental factors such as seasonality or minimum temperature if the traits investigated at a phenotypic level in adults and offspring are driven by the selective pressures of shorter season length and harsher winter temperatures.

Understanding the patterns and processes of adaptation to novel environments over the course of a range expansion in *A. bruennichi* will provide insight into what we can expect as organisms increasingly shift their ranges and encounter novel environments in the course of global climate change. Our approach, which considers different fitness components at different life stages and combines inferences from physiology, morphology, and genomics, provides an example of the kind of integrative studies which will be required for a comprehensive understanding of organismal responses to climate change.

## Material and Methods

### Population samples

We collected adult *A. bruennichi* females from southern France to Estonia from July 29^th^-August 10^th^, 2018, spanning a gradient of 15 degrees of latitude. Dense sampling was done along a transect from southwest to northeast Germany from 23 sites. We sampled sites with as few as three individuals but aimed to collect 10-20 individuals per site (10 or more were collected in 78% of cases). We further collected 120 females from four sites in southwest France, which we call the “core” region and 124 females from three sites in Southern Estonia which we consider the “edge” region. Collecting was timed according to the mating season, when the females had mated (indicated by the lack of males, which are cannibalized during mating) but before egg laying (indicated by the lack of egg sacs in the field).

For the population genetic analysis, we used the adult spiders from the transect through Germany and 10 from each site from the core and edge regions. To fill gaps, we used DNA extracts from samples collected in France, Poland, Lithuania and Latvia by Krehenwinkel and colleagues (list given in Krehenwinkel & Tautz, 2013).

For the common garden experiment, we housed 80 mated female spiders from the core region and 94 from the edge region under common conditions to produce egg sacs. Their egg sacs were then overwintered under native and reciprocal winter conditions for the edge and core region (see below). Collecting locations are visualized in **Figure 1**. GPS coordinates and sample sizes per site can be found in **Supplementary Table S1**.

**Figure 1.**
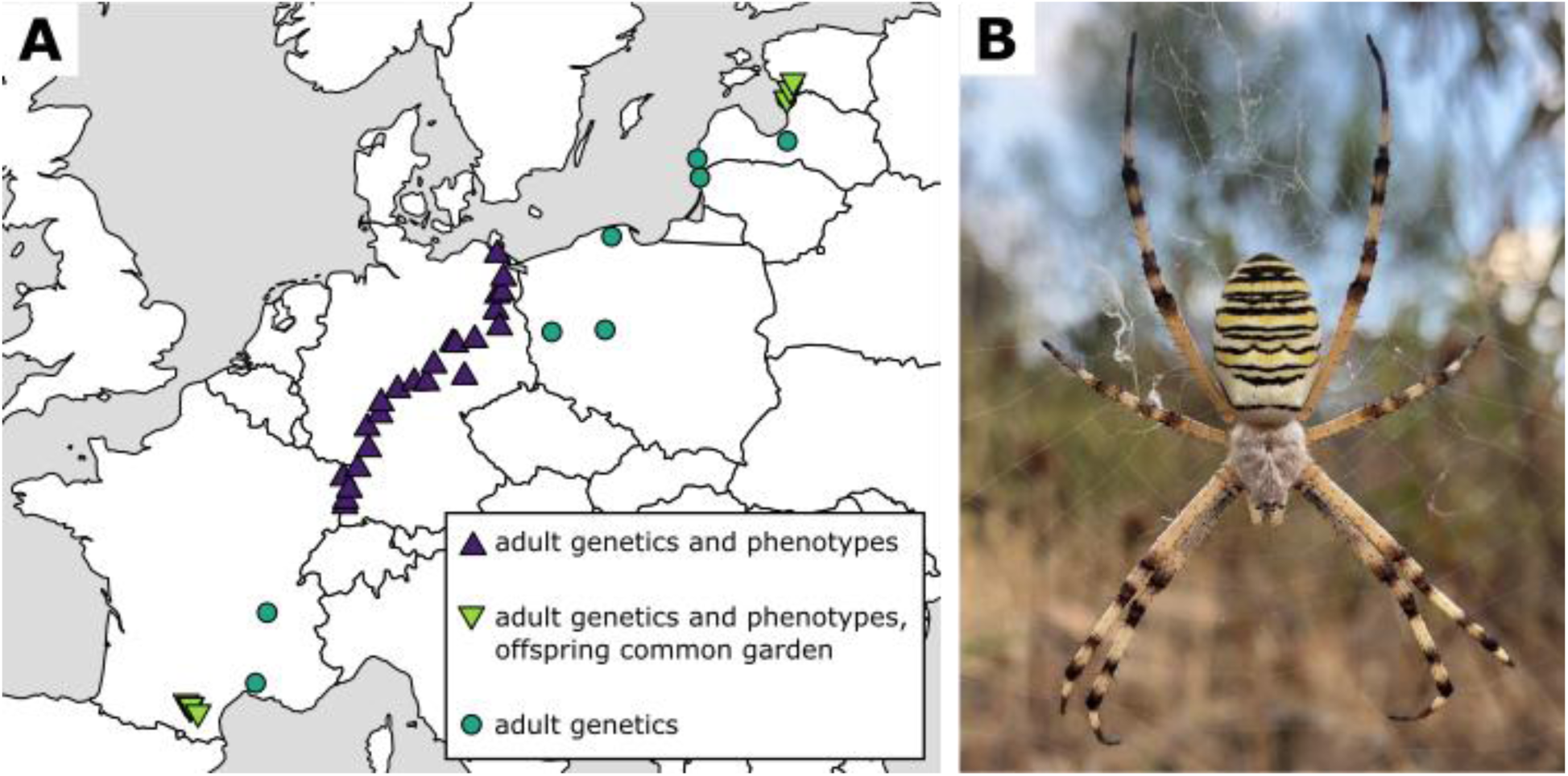
(A) Collecting sites of *Argiope bruennichi* females and utilization for assessment of phenotypic life history traits, population genetics and common garden experiment of overwintering offspring. Each point represents a collecting locality. Samples indicated with triangles refer to females and their offspring collected in 2018 that were used for phenotypic assessment (all) and for the reciprocal common garden winter treatments (extremes of the distribution). Some older, preserved samples (circles) were added for population genetics. (B) Female *Argiope bruennichi* spider in orb web. Photograph taken in Loulé (Faro, Portugal) by Monica M. Sheffer.

### Phenology, female phenotypes and life history traits across latitude

Qualitative data on phenology were collected over the course of several years from 2015 until 2022 for the core and edge sites by recording an estimate of the abundance of adult females and subadult females, and the presence/absence of males and egg sacs in the field. For the females collected across latitude in 2018, we characterized the pigmentation of the dorsal side of their opisthosoma by taking a photograph of each female on the day of collecting. Details of the method for photographing and characterizing pigmentation are found in **Supplementary Text S1**. After collection, we brought the female spiders into a climate-controlled chamber at the University of Greifswald, Germany, where they oviposited under shared conditions (80 % relative humidity, 26°C day temperature, 20°C night temperature, with 12 hours of full day, 10 hours of full night, and one hour to transition between the day and night temperatures): shared conditions were chosen due to similar temperature conditions at the collecting sites in Estonia and France in 2018. The spiders were stored in overturned plastic containers (height: 13 cm, diameter: 12 cm) with a hole filled with cotton on the top for ventilation; additional humidity was provided by spraying the cotton with water every evening. We fed the spiders with adults or larvae of *Lucilia caesar* and *Calliphora vomitoria* flies twice a week. Egg sacs were laid between August 4, 2018 and September 19, 2018. We calculated the days between collection and oviposition and referred to this as “oviposition latency.” The adult females were narcotized after oviposition using CO_2_, preserved in pure ethanol and stored at −80°C. We measured female body size by taking a photograph of the first leg on the right side from each female using a Zeiss Discovery V20 Microscope in combination with a Zeiss AxioCam MRc camera with a 0.63X objective (Zeiss AG, Oberkochen, Germany). From the photos, we measured tibia-patella length as a commonly used proxy for body size in spiders (e.g. Schneider et al., 2006; Uhl & Vollrath, 2000).

*A. bruennichi* females produce intricately woven egg sacs. They are flask-shaped silken containers with an inner chamber hosting the egg mass and an outer chamber in which the spiderlings overwinter (Becker, 1981). Egg sacs were transferred the day following oviposition into plastic boxes (5 x 5 x 3.5 cm) with mesh on two sides. They were then moved into climate cabinets programmed with the same day/night conditions under which they were laid. Eggs develop and spiderlings hatch from the eggs and molt within about four weeks following oviposition and remain in the egg sac as “spiderlings” until they emerge in the following spring (Welke & Schneider, 2012). We refer to the eggs in each egg sac as a “clutch.” Egg sacs from Germany were opened approximately two months after oviposition (64 +/- 13 days) to assess female reproductive success and spiderling size before winter. We calculated the clutch size of the egg sac as the sum of eggs, dead spiderlings, and living spiderlings. Hatching success was calculated as the sum of dead and living spiderlings (“hatched” spiderlings) divided by the total clutch size. Offspring body size was measured as tibia-patella length of the first leg, as for adult females. Egg sacs from France and Estonia were opened later, 8 months after oviposition, as they were subjected to winter treatments in the reciprocal common garden experiment (see below).

To test if the traits adult body size, pigmentation, oviposition latency, clutch size, hatching success and offspring size varied with latitude, we used generalized linear mixed models in R, using the package ‘glmmTMB’ (Brooks et al., 2017). The traits were used as response variables, with latitude (°N) as an explanatory variable. We included maternal body size as an additional explanatory variable for oviposition latency. For clutch-related measurements, we included both maternal body size and oviposition latency as additional explanatory variables. For offspring body size, we also included clutch size as an explanatory variable. When multiple continuous explanatory variables were included, we centered and scaled them in order to allow for interpretation of effects on the same scale (Schielzeth, 2010). Sampling site was included as a random effect. All models fit linear assumptions, assessed with DHARMa (Hartig, 2021). Details of the distribution families and link functions for each model are given in **Supplementary Text S1**. Where possible, we used the Nakagawa & Schielzeth method (Nakagawa & Schielzeth, 2013) to calculate a pseudo-R^2^, implemented in the ‘performance’ R package (Lüdecke et al., 2021), to assess how much of the variation in the data was explained by the full model (conditional pseudo-R^2^), and by the model without random effects (marginal pseudo-R^2^). The Nakagawa & Schielzeth method is not reliable for betabinomial models, thus we used a crude calculation to understand the goodness of fit for our betabinomial models (pigmentation and hatching success); precisely, we calculated the squared correlation of the predicted values with the observed values of the response variable. This does not provide a comparison of conditional and marginal R^2^, thus for betabinomial models we only report the conditional R^2^.

### Reciprocal common garden experiment

#### Offspring phenotypes and cold tolerance

On October 1^st^, 2018, 119 egg sacs from the core and edge regions were assigned to warm or cold winter treatments by random selection, while ensuring approximately equal sample sizes per origin and treatment: core/warm (31 egg sacs), core/cold (28), edge/cold (30) and edge/warm (30). Warm and cold winter treatments were simulated in climate cabinets, the “warm” winter treatment corresponding to the natural conditions of the French sites (climate cabinet: Panasonic MLR-352H, Ewald Innovationstechnik GmbH, Bad Nenndorf, Germany), and the “cold” winter treatment corresponding to the natural conditions of the Estonian sites (climate cabinet: Percival LT-36VL, CLF PlantClimatics GmbH, Wertingen, Germany). For further details on the “cold” and “warm” winter regimes, see **Supplementary Figure S3.**

The egg sacs were opened eight months (243 +/- 1 days) after oviposition between April 4 and May 19, 2019. Most of the egg sacs contained live spiderlings: core/warm (28 egg sacs), core/cold (26), edge/cold (26) and edge/warm (30). We established clutch size and hatching success as described above. Overwintering survival was calculated as the number of living spiderlings divided by the total number of hatched spiderlings. After counting the spiderlings, we took the mass of 15 randomly chosen individual spiderlings per clutch using a Sartorius ME5 microbalance (Sartorius AG, Goettingen, Germany). After weighing, spiderling body size was measured on legs as described above. It can be assumed that body size remains stable over winter, as the leg podomeres are sclerotized body parts and the spiderlings do not molt further until they emerge from the egg sac in spring.

Spiderlings from the four treatments were subjected to cold tolerance measurements. Lower lethal temperature: we measured proportional survivorship across temperatures ranging from −10 to −32°C (20 spiderlings per egg sac). Supercooling point: the temperature at which an organism’s body water freezes (8 spiderlings per egg sac). Chill coma recovery time: the time it takes for an organism to resume coordinated movement after chilling injury (10 spiderlings per egg sac). In some families with low survival, small clutch size, or low hatching success, slightly fewer spiderlings were used in some cold tolerance experiments. Sample sizes for each measurement are found in the results section (**Table 2**). Details of the methods, protocols and precise statistical approach for each measurement of the reciprocal common garden experiment (phenotypes and cold tolerance traits) are given in **Supplementary Text S2.** In a small separate experiment, we determined that these spiders have a freeze-avoidant cold tolerance strategy, meaning they survive up to the point of freezing, and experience freezing-induced mortality (**Supplementary Text S3**).

#### Metabolomics

For a subset of the egg sacs which had very high offspring numbers (core/warm: 7 egg sacs, core/cold: 7; edge/cold: 7 and edge/warm: 5), we performed an exploratory analysis of spiderlings’ metabolome using gas chromatography mass spectrometry (GC-MS). Briefly, intracellular metabolites were extracted from ∼50mg of spiderlings pooled by family. The spiderlings were stored at −80°C immediately after opening the egg sac. Homogenizing of the samples was done using three rounds of bead beating with a FastPrep homogenizer (MP Biomedicals, Irvine, California) and Lysing Matrix E (MP Biomedicals, Irvine, California), 2000 µl of dichloromethane, 6000 µl of methanol, and 200 µl of an internal GC-MS standard (20 nM of N,N-dimethyl-phenylalanine, Sigma-Aldrich). Samples were analyzed by GC-MS as described by Liebeke et al. (2008). Lyophilized samples were derivatized with 40 μl methoxyaminehydrochloride for 90 min at 37°C, mixed with 80 μl N-methyl-N-trimethylsilyltrifluoroacetamide and incubated for 30 min at 37°C. Subsequently, 2 µl of each sample were injected into an Agilent 6890N GC system with an SSl-injector [Split 1:25 at 250°C; inlet split flow: 20 ml/min; carrier gas: helium 1 ml/min (60 kPa) at 110°C; pressure rise: 6 kPa/min] coupled to an Agilent®5973 Network MSD mass selective detector (Agilent Technologies, Santa Clara, Ca, USA) operated in electron ionization mode with an ionization energy of 70 eV. Chromatographic separation was achieved using a 30-m DB-5MS column (30 m x 0,25 mm x 0,25 µm; Agilent Technologies, Santa Clara, Ca, USA) using an oven program comprising the following steps: (1) an initial temperature hold at 70°C for 1 min, (2) stepwise heating with 1.5°C/min up to 76°C (3) stepwise heating with 5°C/min up to 220°C, (3) stepwise heating with 20°C/min up to 325°C, (4) a hold at 325 °C for 8 min. Finally, analytes were transferred to the mass selective detector via the transfer line at 280°C and full scans were performed from 50 to 550 m/z at a scan rate of 2.74 scans per second and a 6 min solvent delay.

The detected compounds were identified by processing the raw GC-MS data with MassHunter version B 8.00 software (Agilent, United States). Retention times and fragmentation patterns of detected metabolites were first aligned to retention times and fragmentation patterns of internal standards, and searched against the NIST 2017 mass spectral database 2.0 d (National Institute of Standards and Technology, Gaithersburg, USA) (Kramida & Ralchenko, 1999) and additionally against an in-house database. Finally, relative concentrations of identified metabolites were calculated based on peak areas of the quantifier ion of each metabolite normalized to the peak areas of the quantifier ion of the internal standard *N,N-*dimethyl-phenylalanine. We compared the relative concentrations of each metabolite according to winter treatment (warm/cold), origin (core/edge), and the interaction of winter treatment and origin using linear models. We additionally included the identity of who performed the extraction as an additive fixed effect to account for differences in extraction efficiency.

### Genomic patterns across latitude

#### Sequencing and data filtration

We extracted genomic DNA from the four left legs of adult female spiders using a plate extraction protocol, prepared dual-indexed libraries using a double-digest restriction-enzyme-associated DNA sequencing (ddRADseq) library preparation protocol (adapted from Peterson et al., 2012). We performed sequencing read processing and SNP filtering, briefly described here, with more detail provided in **Supplementary Text S4**. We removed adapter sequences using Trimmomatic (Bolger et al., 2014). Next, we mapped the reads of each sample onto the current reference genome assembly for this species (GCA_015342795.1, Sheffer *et al*., 2021) using BWA (Li, 2013) and then filtered the SNP data, retaining SNPs for downstream analyses if they had a minimum coverage of 3x per individual, no more than 40**%** of missing data for individual sites, and a maximum coverage of 240x per individual using SNPcleaner (Fumagalli et al., 2014). The cutoff for maximum coverage is about 10 times higher than the average coverage per individual, and helps to avoid SNP calls that fall in highly repetitive regions of the genome. We used a custom pipeline (Hoff, unpublished; see Data Availability section) to further filter the SNPs based on their position inside/outside of exons and introns. Intergenic SNPs, which did not fall within introns or exons, were used for population genomic analyses that call for neutral loci. We will refer to these as “neutral SNPs” henceforth. For analyses which included all data, including those SNPs falling within genes, we will refer to “all SNPs.” To account for uncertainty in SNP and genotype calls based on allele counts, which might introduce noise or bias into downstream analyses (Johnson & Slatkin, 2008; Lynch, 2008), we used an empirical Bayesian framework, implemented in ANGSD (<u>a</u>nalysis of <u>n</u>ext-<u>g</u>eneration <u>s</u>equencing <u>d</u>ata; Korneliussen et al., 2014) to calculate genotype likelihoods instead of generating SNP genotype calls whenever possible.

#### Population genomics

Using the neutral SNPs, we characterized the population genetic structure and assessed demographic processes, briefly described here. Greater detail on the precise methods and tools used can be found in **Supplementary Text S4**. We performed a principal component analysis (PCA) as implemented in ngsTools “PCAngsd” (Fumagalli et al., 2013), and calculated individual admixture proportions for assignment of sites into genetic clusters using ngsAdmix (Skotte et al., 2013). For insight into demographics of our populations, we calculated per-individual inbreeding coefficients and pairwise relatedness, and pairwise F_ST_ between each sampling site (Vieira et al., 2013, 2016).

#### Genome-wide test for environmental adaptation

To test for environmental adaptation we used genotype-environment association tests (e.g. Ahrens et al., 2018; Dudaniec et al., 2018; Hoban et al., 2016). We tested the association of genotypes with climate data that were extracted for all collecting sites along the transect. We used the 19 standard bioclimatic variables from WorldClim at 30 second (∼ 1km^2^) resolution (Fick & Hijmans, 2017). To reduce the number of environmental variables in downstream analyses, we summarized the 19 variables into fewer dimensions using a principal component analysis (PCA) using the R package ‘FactoExtra’ (Kassambara & Mundt, 2020) (**Supplementary Text S5**), following the recommendation of Hoban et al. (2016), but also visually assessed the individual climatic variables across our sampling sites (**Supplementary Figure S10**). We performed the association tests in ANGSD using the **-**doAsso method, which uses a generalized linear framework to associate genotype likelihoods with binary, count, or continuous variables (Skotte et al., 2012). This test calculates a likelihood ratio test (LRT) score, and we use a p-value cut-off of 0.0005 based on a chi-square distribution, corresponding to an LRT of 12.116 as our significance threshold.

## Results

### Phenology, female phenotypes and life history traits across latitude

We assessed how phenology and adult female phenotypic and life history traits (opisthosoma pigmentation, body size, fecundity, hatching success and offspring size) vary with latitude (**Figure 2**, **Table 1, Supplementary Figure S2**). Phenology differs between the core and edge region in that females of the edge region were mature in the first half of August, always about two weeks earlier compared to females of the core region. This pattern was found repeatedly in the years 2015 to 2021. Further, females in the edge region are more synchronized in their reproductive development, with no males and subadult females or only singletons being found already at the beginning of August, whereas the reproductive season was more spread out in the core region, with mostly adult females but also adult males, subadult males and subadult females being found later in the summer. Opisthosoma pigmentation of the females did not significantly vary with latitude (**Figure 2A**, **Table 1**). Female body size decreased significantly with increasing latitude (**Figure 2B**, **Table 1**). There was no significant effect of latitude on female fecundity (measured as clutch size of the first egg sac) (**Figure 2C**, **Table 1**). Generally, larger females produce larger clutches independent of origin (**Table 1**). Oviposition latency was shorter for spiders from increasingly high latitudes (**Table 1, Supplementary Figure S2**). Hatching success, as the proportion of spiderlings hatched out of the total eggs laid per egg sac, had a mean of 0.964 (±0.112) and did not vary significantly with latitude (**Figure 2D**, **Table 1**), nor did the size of the spiderlings (mean ± standard deviation: 559.523 ± 39.212 µm) (**Table 1, Supplementary Figure S2**).

**Figure 2:**
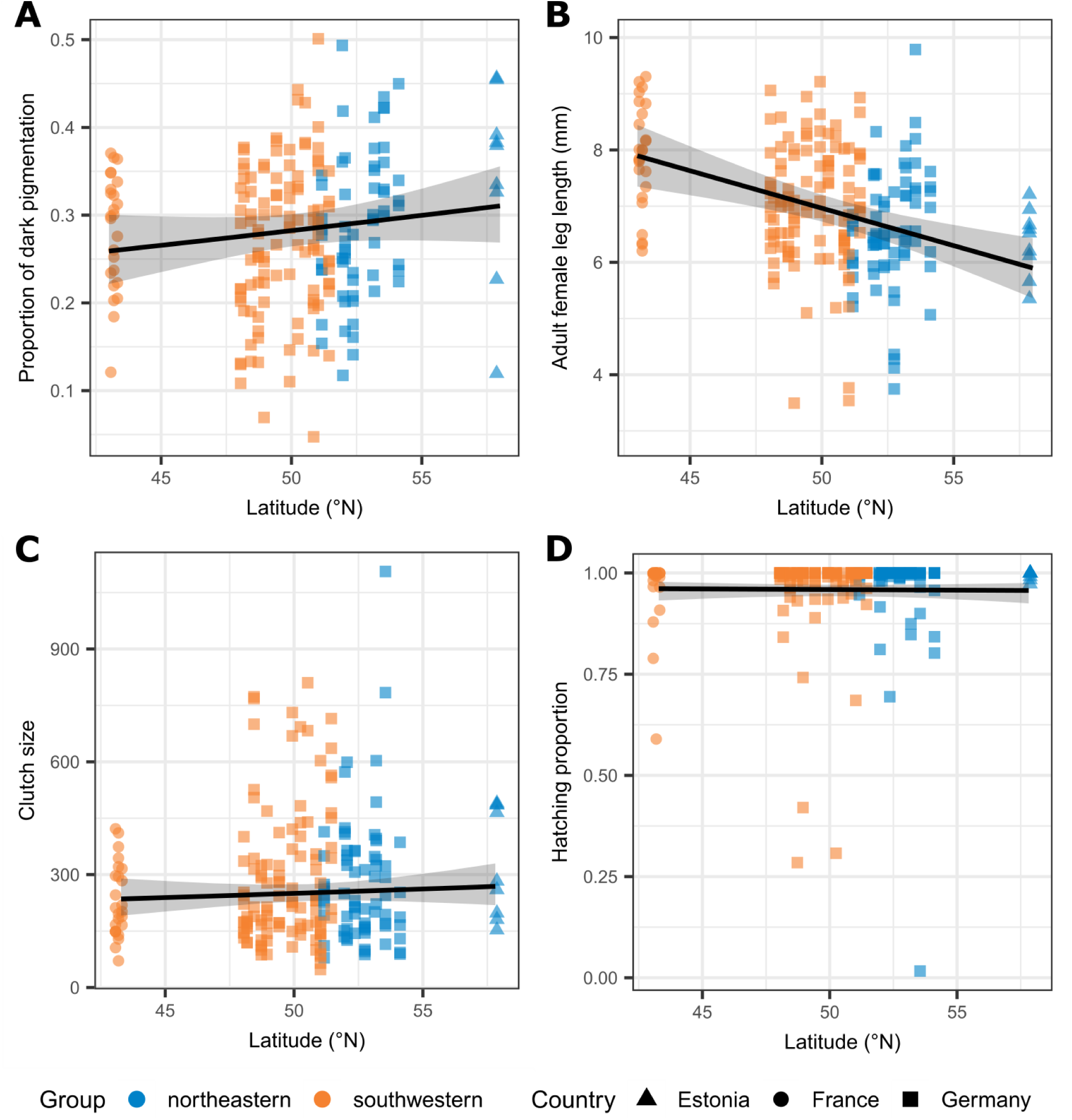
Phenotypic traits of adult *A. bruennichi* females according to latitude. Points are colored according to their assignment to “southwestern” (orange) or “northeastern” (blue) genetic clusters (see population genetic results, Figure 4) and shaped according to the sampling country. Model estimates are plotted as black lines with their 95% confidence intervals as transparent gray ribbons. (**A**) Opisthosoma pigmentation, as the proportion of dark to light pixels. (**B**) Adult female tibia-patella length of the first leg (in mm), a common body size proxy in spiders. (**C**) Number of offspring (spiderlings) per clutch (egg sac). (**D**) Proportion of spiderlings hatching per egg sac.

**Table 1.** Model output for phenotypic traits of adult *A. bruennichi* females according to latitude. Estimates are reported on the latent scale, with 95% confidence intervals. Fixed effects are considered significant if their confidence interval does not cross zero; these are indicated in **bold text.**

| Response Variable | Sample size | Pseudo-R<sup>2</sup> (cond.) | Pseudo-R<sup>2</sup> (marg.) | Fixed effects | 2.5% | 97.5% | Estimate |
| --- | --- | --- | --- | --- | --- | --- | --- |
| Opisthosoma pigmentation | 262 | 0.319 | NA | Latitude | -0.0076 | 0.0414 | 0.0169 |
| Adult female body size | 273 | 0.381 | 0.172 | **Latitude** | -198.4733 | -68.0019 | -133.2376 |
| Oviposition latency | 273 | 0.156 | 0.061 | **Latitude\*** | -0.2935 | -0.0466 | -0.1701 |
|  |  |  |  | Mother's body size* | -0.1091 | 0.0943 | -0.0074 |
| Clutch size | 273 | 0.627 | 0.490 | Latitude* | -0.0603 | 0.1265 | 0.0331 |
|  |  |  |  | **Mother's body size\*** | 0.1746 | 0.2884 | 0.2315 |
|  |  |  |  | **Oviposition latency\*** | -0.3837 | -0.2818 | -0.332 |
| Hatching success | 273 | 0.038 | NA | Latitude* | -0.2636 | 0.2074 | -0.0281 |
|  |  |  |  | Mother's body size* | -0.4421 | 0.0505 | -0.1958 |
|  |  |  |  | Oviposition latency* | -0.4135 | 0.0404 | -0.1866 |
| Offspring body size | 2999 | 0.801 | 0.016 | Latitude* | -5.9402 | 15.2234 | 4.6416 |
|  |  |  |  | Mother's body size* | -5.0192 | 7.4859 | 1.2334 |
|  |  |  |  | Oviposition latency* | -8.6031 | 3.5255 | -2.5388 |
|  |  |  |  | Clutch size* | -5.2738 | 6.5996 | 0.6629 |

\*centered and scaled in the model

### Reciprocal common garden experiment with overwintering spiderlings

#### Overwintering survival

We found no significant differences in survival over winter between spiderlings from core and edge origins. There was a strong effect of decreased survival in the cold winter treatment for spiders from both origins (estimated mean survival proportions: core/warm: 0.84 (95% confidence interval (CI) 0.76-0.90), core/cold: 0.58 (95%CI 0.46-0.70), edge/warm: 0.81 (95%CI 0.71-0.88), edge/cold: 0.61 (95%CI 0.49-0.72)), and slightly decreased survival with increasing maternal body size. The model had a pseudo-R^2^ of 0.319 (**Figure 3A**, **Table 2**).

**Figure 3:**
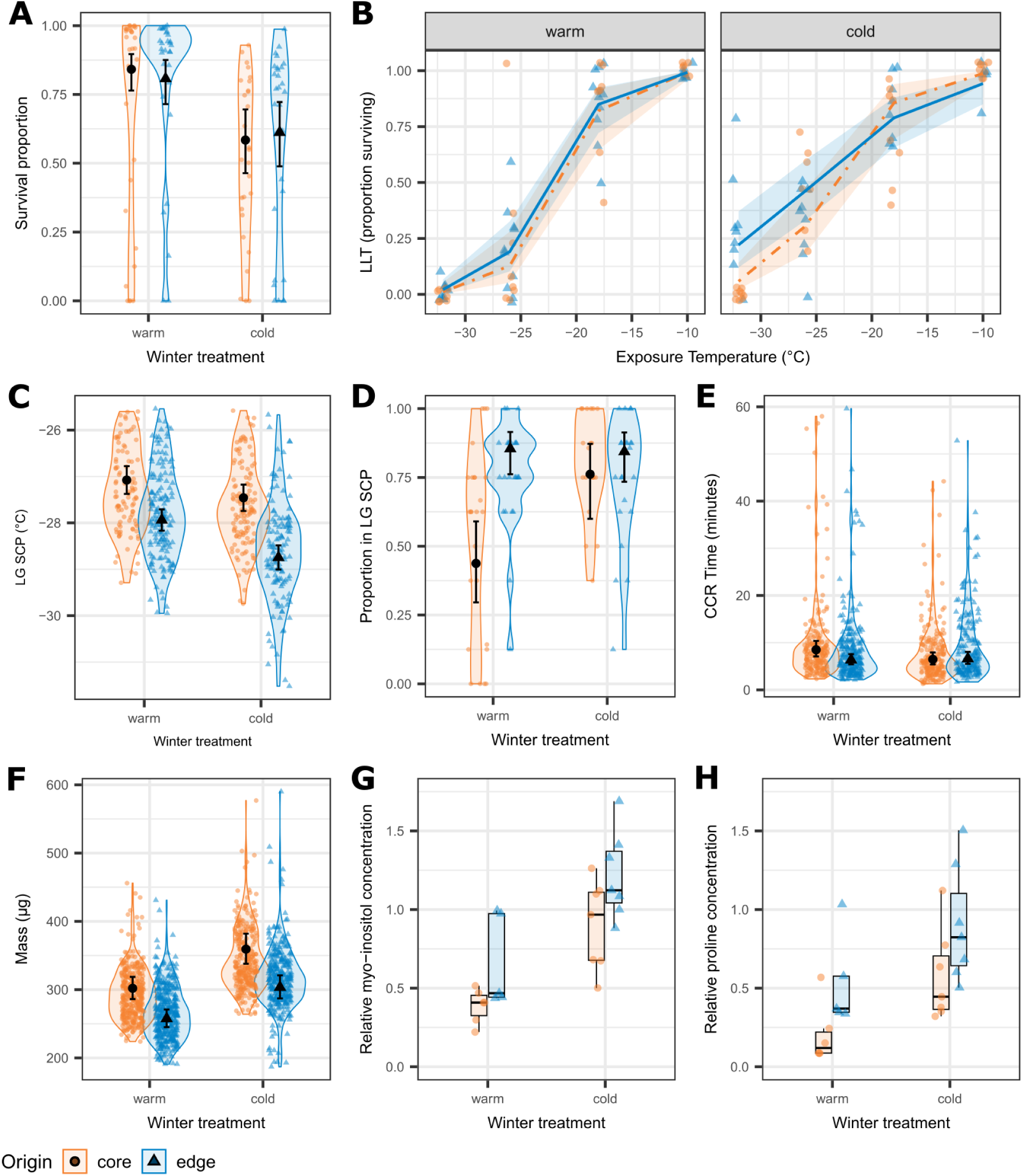
Reciprocal common garden experiment: survival, cold tolerance traits, mass and metabolite concentrations of spiderlings according to origin and winter treatment. (**A**) Overwintering survival of spiderlings according to winter treatment (warm or cold) and origin (core or edge of the range). (**B**) Lower lethal temperature (LLT) according to exposure temperature, winter treatment and origin. (**C**) Low-group (LG) supercooling points (SCPs) according to winter treatment and origin. (**D**) Within a family (spiderlings from an egg sac) the proportion of supercooling points (SCPs) that are in the “low group” (LG). A high proportion indicates that most of the individuals from that family had LG SCPs, a low proportion means that most had high group SCPs. (**E**) Chill coma recovery time in minutes. (**F**) Mass of spiderlings in micrograms. Black points with error bars (A, C, D, E, F) or lines with ribbons (B) represent model predictions flanked with their 95% confidence intervals. Model predictions are back-transformed to the observed scale. Semi-transparent points represent the raw data, horizontally jittered to allow visualization of overlapping points. The color, shape, and line type all represent the origin. (**G**) Relative concentrations of myo-inositol, a cyclic polyol, according to winter treatment and origin. (**H**) Relative concentrations of proline, an amino acid, according to winter treatment and origin. For (G) and (H), single points represent pooled spiderlings from a single egg sac extracted as a group, with color and shape indicating their population of origin (core or edge), with horizontal jittering to show overlapping points more clearly. Boxes represent the interquartile range, also colored by population of origin, thick lines represent medians, and whiskers represent 1.5 times the interquartile range.

**Table 2:**
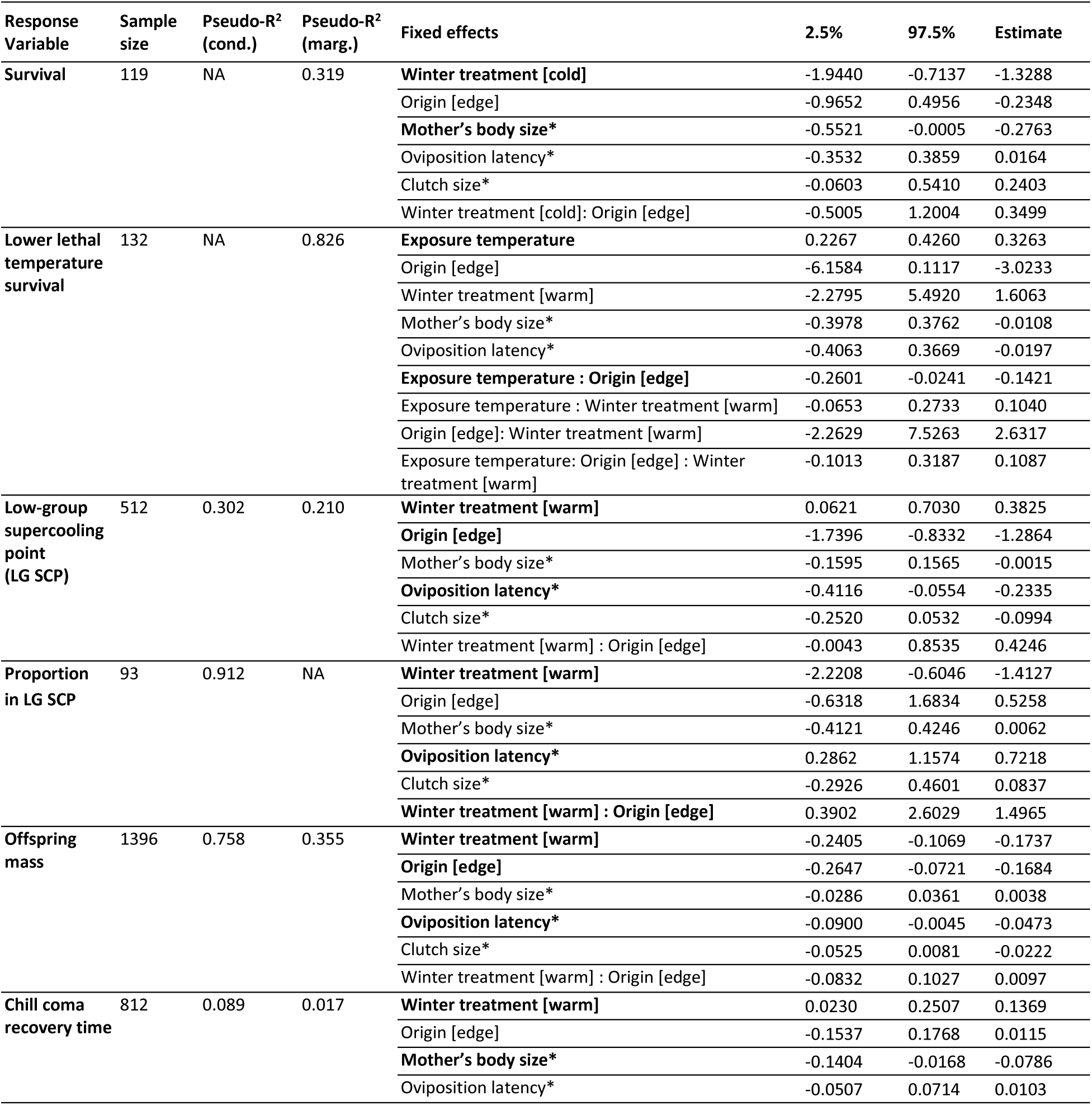
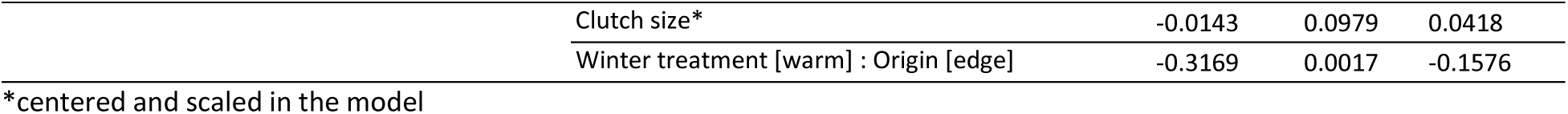
Reciprocal common garden experiment: model output for offspring traits. Estimates are reported on the latent scale, with 95% confidence intervals. Fixed effects are considered significant if their confidence interval does not cross zero; these are indicated in **bold text.** Fixed effects separated by a colon indicate an interaction of those effects. Reference level of fixed effects is given in [square brackets].

#### Lower lethal temperature

In general, survival decreased with colder exposure temperatures. The effect of origin was only significant in combination with exposure temperature. At the coldest exposure temperatures (−26°C and −32°C), spiders from the edge of the range, overwintered under cold conditions, survive at slightly higher proportions compared to spiders from the core (estimated mean survival proportions at −32°C: core/warm: 0.01 (95%CI 0.00-0.05), core/cold: 0.06 (95%CI 0.02-0.14), edge/warm: 0.02 (95%CI 0.01-0.07), edge/cold: 0.22 (95%CI 0.12-0.37), and at −26°C: core/warm: 0.13 (95%CI 0.05-0.28), core/cold: 0.30 (95%CI 0.18-0.47), edge/warm: 0.19 (95%CI 0.10-0.33), edge/cold: 0.46 (95%CI 0.33-0.60)). The pseudo-R^2^ for the model was 0.826 (**Figure 3B**, **Table 2**).

#### Supercooling points

We analyzed the data on supercooling points (SCPs) of spiderlings from the four treatment groups with two models, due to multimodality of the raw data: firstly, we analyzed the “low group” (LG) SCPs as a continuous variable. SCPs were classified into the LG if they were below −25.5°C, accounting for 77.5% of the data. Secondly, we analyzed the per-clutch ratio of LG to “high group” (HG) SCPs (SCPs higher than −25.5°C, **see Supplementary Text S2, Figure S5**) to test for differences in supercooling strategy among treatment groups. For the LG, our model showed a significantly lower SCP for spiderlings originating from the edge, and higher SCP for both origins under warm conditions (estimated mean SCP: core/warm: −27.08°C (95%CI −27.38 to −26.78), core/cold: −27.46°C (95%CI −27.74 to −27.17), edge/warm: −27.94°C (95%CI −28.17 to −27.71), edge/cold: −28.75°C (95%CI −29.01 to −28.49) (**Figure 3C**). The SCP was slightly lower with increasing oviposition latency. The LG SCP model had a conditional pseudo-R^2^ of 0.302 and a marginal pseudo-R^2^ of 0.210 (**Figure 3C**, **Table 2**). The model comparing the ratio of LG:HG SCPs had a pseudo-R^2^ of 0.912. There were more HG SCPs in the warm winter treatment and with longer oviposition latency (**Table 2**). There was a strongly significant positive effect of the interaction of winter treatment and origin: more than 50% of spiderlings with a core origin under warm conditions have high group supercooling points, while edge spiderlings maintain a strong majority of low group supercooling points in both winter treatments (estimated mean proportion in LG: core/warm: 0.44 (95%CI 0.30-0.59), core/cold: 0.76 (95%CI 0.60-0.87), edge/warm: 0.85 (95%CI 0.76-0.92), edge/cold: 0.84 (95%CI 0.73-0.91), **Figure 3D**, **Table 2**). In an additional experiment, we determined if spiderlings indeed survive up to the point of freezing, and thus experience freezing-induced mortality. These tests suggest that spiderlings employ a freeze-avoidant cold tolerance strategy (**Supplementary Text S3, Figure S6**).

#### Chill coma recovery time

We found no significant differences in chill coma recovery time between spiderlings from different origins (core versus edge). Chill coma recovery time was shorter in the cold winter treatment for spiders from both regions (estimated mean chill coma recovery time in minutes: core/warm: 8.51 (95%CI 7.09-10.38), core/cold: 6.47 (95%CI 5.39-7.92), edge/warm: 6.36 (95%CI 5.42-7.57), edge/cold: 6.61 (95%CI 5.52-8.07)). Chill coma recovery time was faster with increasing maternal body size. The model had a very low conditional pseudo-R^2^ of 0.089 and marginal pseudo-R^2^ of 0.017 (**Figure 3E**, **Table 2**).

#### Body mass, size and condition

We found significant differences in mass between origins and winter treatments, with edge spiderlings consistently lighter than those from the core, and an overall higher mass under cold winter conditions (estimated mean mass in micrograms: core/warm: 299.92 (95%CI 285.90-314.63), core/cold: 356.16 (95%CI 337.42-375.94), edge/warm: 259.14 (95%CI 247.01-271.86), edge/cold: 311.51 (95%CI 296.37-327.43). We also observed a small but significant decrease in mass with longer oviposition latency. The conditional pseudo-R^2^ of the model was 0.758, marginal pseudo-R^2^ 0.355 (**Figure 3F**, **Table 2**). The results for body condition are very similar to the results for mass, although leg length differences are not significantly different (**Supplementary Figure S4**).

#### Metabolomics

GC-MS analysis identified 46 metabolites in our samples (**Supplementary Table S2**). Of these, 10 showed significantly greater relative concentrations in the cold treatment and/or in edge spiderlings, which we focus on as potential effects of cold temperatures, and/or mechanisms to enhance cold tolerance: five amino acids (valine, leucine, isoleucine, proline, lysine), three tricarboxylic acid (TCA) cycle intermediates (citrate, itaconate, aconitate), one glycolysis intermediate (3-phosphoglycerate) and one polyol (myo-inositol) (**Figure 3G-H**, **Table 3**).

**Table 3:** Reciprocal common garden experiment: model output for metabolite concentrations. Estimates are reported on the latent scale, along with the standard error (“SE”), sum of squares (“Sum Sq”), F value, and p-value (“Pr(>F)”). Fixed effects are considered significant if the p-value is below 0.05; these are indicated in **bold text.** Fixed effects separated by a colon indicate an interaction of those effects. We included the identity of the person performing the metabolite as “extractor” in the model to account for differences between people performing the extractions. Reference level of fixed effects is given in [square brackets].

| Response Variable | Compound class | Fixed effects | Estimate | SE | Sum Sq | F value | Pr(>F) |
| --- | --- | --- | --- | --- | --- | --- | --- |
| Valine | Amino acid | **Winter treatment [cold]** | -0.530 | 0.229 | 1.994 | 11.758 | 0.003 |
|  |  | **Origin [edge]** | 0.389 | 0.220 | 0.956 | 5.637 | 0.028 |
|  |  | Extractor | -0.331 | 0.172 | 0.615 | 3.624 | 0.071 |
|  |  | Winter treatment [cold] : Origin [edge] | -0.094 | 0.337 | 0.013 | 0.078 | 0.783 |
| Leucine | Amino acid | **Winter treatment [cold]** | -0.879 | 0.251 | 3.621 | 17.896 | 0.000 |
|  |  | Origin [edge] | 0.012 | 0.240 | 0.114 | 0.564 | 0.461 |
|  |  | Extractor | -0.204 | 0.188 | 0.287 | 1.417 | 0.248 |
|  |  | Winter treatment [cold] : Origin [edge] | 0.220 | 0.368 | 0.072 | 0.358 | 0.556 |
| Isoleucine | Amino acid | **Winter treatment [cold]** | -0.758 | 0.218 | 2.208 | 14.357 | 0.001 |
|  |  | Origin [edge] | -0.056 | 0.210 | 0.068 | 0.445 | 0.513 |
|  |  | Extractor | -0.153 | 0.164 | 0.186 | 1.209 | 0.285 |
|  |  | Winter treatment [cold] : Origin [edge] | 0.322 | 0.321 | 0.155 | 1.011 | 0.327 |
| Proline | Amino acid | **Winter treatment [cold]** | -0.356 | 0.157 | 0.916 | 11.584 | 0.003 |
|  |  | **Origin [edge]** | 0.328 | 0.150 | 0.670 | 8.474 | 0.009 |
|  |  | Extractor | -0.219 | 0.118 | 0.268 | 3.387 | 0.081 |
|  |  | Winter treatment [cold] : Origin [edge] | -0.065 | 0.230 | 0.006 | 0.079 | 0.782 |
| Lysine | Amino acid | **Winter treatment [cold]** | -0.621 | 0.163 | 1.357 | 15.911 | 0.001 |
|  |  | Origin [edge] | -0.054 | 0.156 | 0.076 | 0.894 | 0.356 |
|  |  | **Extractor** | -0.300 | 0.122 | 0.608 | 7.130 | 0.015 |
|  |  | Winter treatment [cold] : Origin [edge] | 0.286 | 0.239 | 0.122 | 1.436 | 0.245 |
| Citrate | TCA cycle intermediate | **Winter treatment [cold]** | -0.993 | 0.158 | 3.437 | 42.768 | 0.000 |
|  |  | Origin [edge] | -0.158 | 0.152 | 0.038 | 0.475 | 0.499 |
|  |  | Extractor | -0.093 | 0.118 | 0.105 | 1.300 | 0.268 |
|  |  | **Winter treatment [cold] : Origin [edge]** | 0.512 | 0.232 | 0.393 | 4.886 | 0.039 |
| Itaconate | TCA cycle intermediate | Winter treatment [cold] | 0.272 | 0.235 | 0.145 | 0.817 | 0.377 |
|  |  | Origin [edge] | 0.710 | 0.225 | 0.718 | 4.047 | 0.058 |
|  |  | Extractor | -0.126 | 0.176 | 0.019 | 0.107 | 0.747 |
|  |  | **Winter treatment [cold] : Origin [edge]** | -0.884 | 0.344 | 1.169 | 6.593 | 0.018 |
| Aconitate | TCA cycle intermediate | **Winter treatment [cold]** | -0.687 | 0.175 | 1.804 | 18.337 | 0.000 |
|  |  | Origin [edge] | -0.190 | 0.168 | 0.053 | 0.536 | 0.473 |
|  |  | **Extractor** | 0.503 | 0.131 | 1.317 | 13.394 | 0.002 |
|  |  | Winter treatment [cold] : Origin [edge] | 0.374 | 0.256 | 0.209 | 2.127 | 0.160 |
| 3-phosphoglycerate | Glycolysis intermediate | Winter treatment [cold] | -0.118 | 0.209 | 1.275 | 9.088 | 0.007 |
|  |  | Origin [edge] | 0.475 | 0.200 | 0.141 | 1.007 | 0.328 |
|  |  | Extractor | 0.294 | 0.157 | 0.700 | 4.988 | 0.037 |
|  |  | Winter treatment [cold] : Origin [edge] | -0.653 | 0.306 | 0.638 | 4.545 | 0.046 |
| Myo-inositol | Cyclic polyol | Winter treatment [cold] | -0.489 | 0.101 | 1.840 | 56.319 | 0.000 |
|  |  | Origin [edge] | 0.316 | 0.097 | 0.556 | 17.008 | 0.001 |
|  |  | Extractor | -0.348 | 0.076 | 0.664 | 20.312 | 0.000 |
|  |  | Winter treatment [cold] : Origin [edge] | -0.144 | 0.148 | 0.031 | 0.951 | 0.341 |

### Population genetic structure and genome-wide evidence for environmental adaptation

To test for genomic differentiation underlying the phenotype-level measurements, we performed a population genetic study of adult females from across the range. After demultiplexing and trimming adapters, ddRAD libraries for the 423 adult females yielded more than 362 million reads, with an average of 815,796 reads per sample (min: 5,296 reads, max: 12,597,258 reads). An average (mean) of 86.89% of reads per sample mapped to the reference genome, with a mean coverage of 4.94X for each of the 13 chromosome-level superscaffolds (Sheffer et al., 2021). Subsequent SNP calling resulted in 22,372 biallelic SNPs. 13,832 of those SNPs did not fall within exons or introns and were used as “neutral” SNPs in population clustering analyses.

In the genetic principal component analysis, PC1 accounted for 17.4% of genetic variation, separating French and southwestern German populations from the northeastern European populations, while PC2 accounted for 3.4% of the variation, separating Estonian populations from the remaining European populations, and PC3 accounted for 2.5% of variation, with some separation of French populations from southwestern German populations, but otherwise little distinction between site groupings (**Figure 4B, Supplementary Figure S9**).

**Figure 4.**
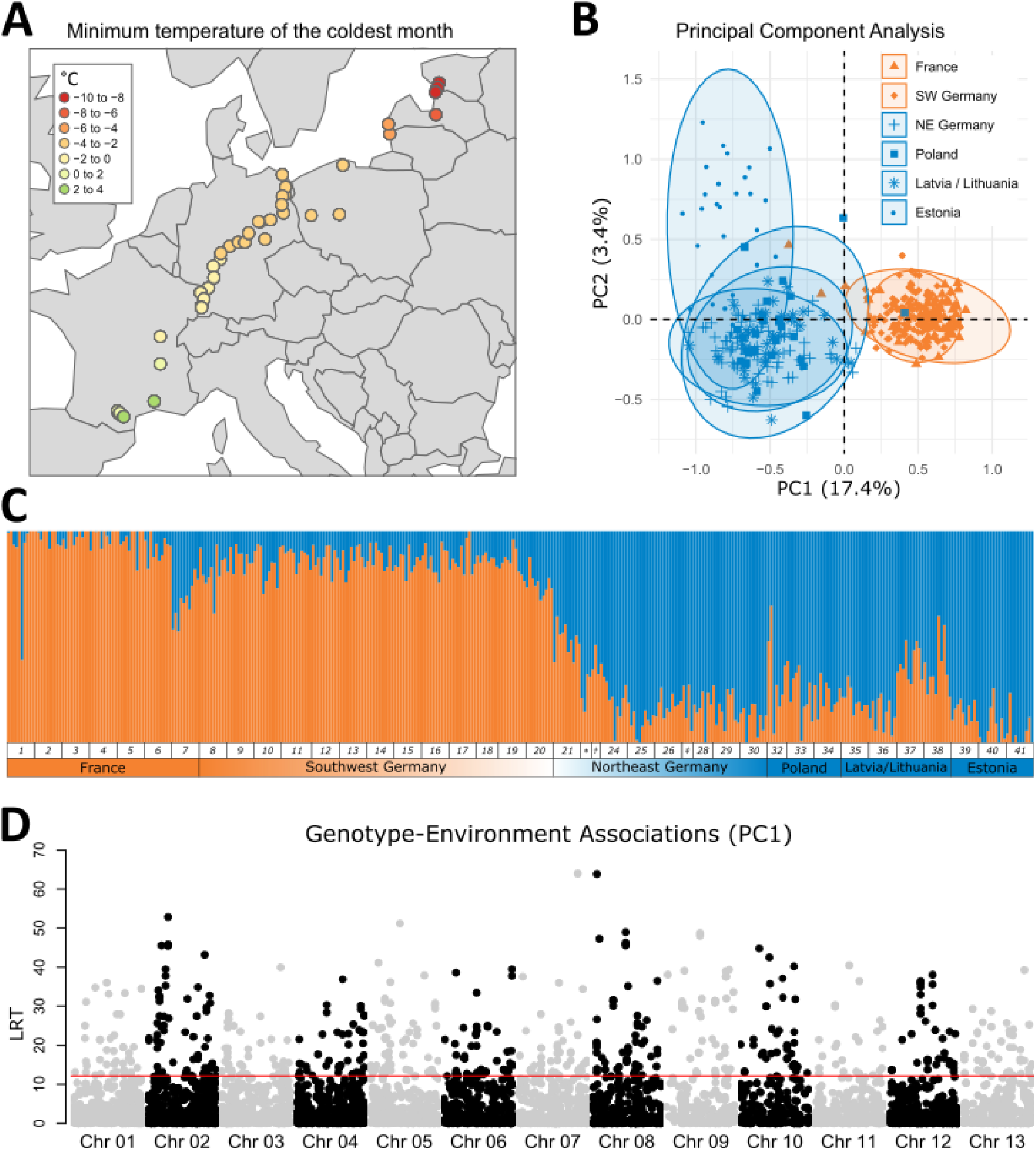
Genetic analyses of *A. bruennichi* (adult females) sampled over a transect from southwestern France to southwestern Estonia (sampling sites: Supplementary Table S1). (A) Sampling sites are represented as points colored according to the bioclimatic variable of minimum temperature of the coldest month (“minTempColdestMonth”), which is averaged over a period of 30 years. (B) Principal component analysis (PCA) using neutral single nucleotide polymorphisms (SNPs). Ellipses represent 95% confidence intervals; points represent individual samples, with shape indicating the sampling country. Points and ellipses are colored according to the site’s “southwest” (orange) or “northeast” (blue) assignment. (C) Stacked bar plots of individual ancestry assignment, using ngsAdmix results for K=2. Colors represent genetic groupings. Numbers below the bars show the sampling sites. For sampling sites with few individuals, symbols were used instead of numbers: * represents population number 22, † represents 23, and ‡ represents 27. (D) Genotype-environment associations of all SNPs with the first axis of a PCA of environmental variables. Individual points show the likelihood ratio test statistic (LRT) that a single SNP is significantly associated with the environment. Grey and black coloration visually distinguishes the SNPs on each chromosome. Chromosome positions are based on alignment to the reference genome. The red line represents a significance threshold of 0.0005, based on a chi-squared distribution LRT value of 12.016.

Using ngsAdmix (Skotte et al., 2013), we calculated individual admixture proportions, with values of *K* ranging from *K*=2 to *K*=20. Here, we present the results for *K*=2, which best shows the transition between southwest and northeast populations (**Figure 4C**) (results for *K=*3 and *K*=4 in **Supplementary Figure S8**). The results show a rapid turnover within Germany: in southwestern Germany, individuals are assigned to a genetic cluster with France, and in the center of Germany there is turnover with individuals being assigned to the cluster containing northeastern European countries. The turnover zone is found in the center of Germany, between sampling sites 19 and 23, with a transition zone of ∼127 kilometers. For visualization purposes, we split the clusters between sites 19 and 20 when plotting with two colors in other figures (e.g. **Figure 4B**, **Figure 2**).

We calculated the levels of inbreeding at every sampling site, to gain insight into the levels of isolation of our populations. The majority of our sites are largely outbreeding, but the populations at the edge of the range in Estonia (population numbers 39, 40, and 41) show an increase in the level of inbreeding with latitude (**Supplementary Figure S7A**). We also calculated the pairwise F_ST_ between all sampling sites, which ranged from 0.025 between a site in Lithuania (site number 35) and a site in Poland (site number 32) to 0.130 between a site in Germany (site number 23) and a site in Estonia (site number 41). F_ST_ was lower among “southwestern” populations (min: 0.030, max: 0.074) than among “northwestern” populations (min 0.025, max: 0.130) (**Supplementary Figure S7B**).

#### Environmental data analysis and genotype-environment associations

We extracted historical climate data for 19 bioclimatic variables (**Supplementary Text S5, Figure S10**) and summarized them into fewer dimensions. PC1 accounted for 51.9% of the variation in the environmental data, and varied almost perfectly with latitude (**Supplementary Figure S10-S11**). PC1 is strongly influenced by temperature variables, especially the mean temperature of the coldest quarter and the minimum temperature of the coldest month (**Figure 4A**). PC2 accounted for 24.4% of variation, and separated sites more according to precipitation than temperature (**Supplementary Figure S12**). PC3 accounted for 11.2% of variation, with the strongest loadings relating to seasonality in temperature and precipitation, as well as the annual temperature range (**Supplementary Figure S13**). An overview of the loadings for each PC is given in **Supplementary Table S3**.

We analyzed genotype-environment associations to identify potential genomic regions underlying adaptation to the environment. Firstly, we looked at associations with the environmental PC1: 589 SNPs had significant associations with this latitudinal temperature variation. We found windows showing significant associations on all chromosomes, with particularly many significant SNPs on chromosome 2, where there is an approximately 30 Mbp-long region showing strong association with the environment (**Figure 4D**). We found 186 significant associations with the environmental PC3, and the same region of chromosome 2 stands out again. For the other environmental principal components, we found fewer significant associations: 84 for PC2, 53 for PC4, and 109 for PC5.

## Discussion

We set out to understand how the wasp spider, *Argiope bruennichi*, has been able to rapidly establish in northeastern Europe, asking whether they have evolved adaptive differences, or been able to cope through phenotypic plasticity. Our results overall suggest that increased seasonality and colder minimum winter temperatures have driven adaptive differentiation to maintain fecundity and enhance low temperature survival in northeastern populations, despite substantial phenotypic plasticity and high gene flow.

Our study incorporates information from across the life cycle of this annual species. At the level of the adult phenotype, our results are suggestive of adaptive responses (through either genetic adaptation or phenotypic plasticity) to the shorter growing season at higher latitudes. Given the longer winters and lower temperatures at the edge of the distribution, we expected individuals of *A. bruennichi* to mature later at the edge compared to the core region, due to restrictions on energy acquisition. Lower temperatures reduce the probability of foraging by impacting activity, as well as through reduced prey availability, both of which should delay maturation. Our observations over several years, however, show that the mating season starts earlier at the edge in *A. bruennichi*, despite these constraints. At range edges, rapid development often seems favored (Willi & Buskirk, 2022), as earlier maturation might be highly selected for in environments that allow only short reproductive seasons, suggesting local adaptation. Rapid development is expected to come with intrinsic costs, such as smaller size and/or reduced fecundity (Willi & Buskirk, 2022). Indeed, in *A. bruennichi*, body size does decrease with increasing latitude in this study and in accordance with previous findings (Krehenwinkel & Tautz, 2013; Wolz et al., 2020), but comes without a concurrent reduction in fecundity: neither clutch size nor hatching success is reduced in northern populations. Lastly, we expected that due to the positive thermoregulatory effects of melanism, and because many ectothermic animals (i.e. reptiles, insects) show darker coloration at higher altitudes and latitudes or when exposed to low temperatures and vice versa (Clusella-Trullas et al., 2008; Guppy, 1986; Heidrich et al., 2018; MacLean, Higgins, et al., 2016; MacLean, Kingsolver, et al., 2016; Martínez-Freiría et al., 2020; Svensson & Waller, 2013; Watt, 1968; Zeuss et al., 2014), northern populations of *A. bruennichi* would be darker in coloration than southern populations. However, we did not find differences in the proportion of dark pigmentation across latitude. There are many plausible explanations for this finding: spiders may perform behavioral thermoregulation by orienting either their dorsal or ventral side toward the sun, as has been documented in some other spider species and within the *Argiope* genus (Robinson & Robinson, 1978; Tolbert, 1979). Alternatively, although the dorsal opisthosoma pattern is a striking feature in this species, the pigmentation of the legs and/or prosoma may be more relevant for thermoregulation, as most of the muscles are situated in these body parts. The dorsal opisthosoma pattern and pigmentation might have evolved under different selection regimes, e.g. to impede the formation of a searching image for predators or to attract insect prey (Bond, 2007; Bush et al., 2008; Croucher et al., 2011), or, temperature may not be as strong a driver as season length or photoperiod.

The adult phenotypes were measured from natural populations, so the differences over latitude that we found may arise from intra- or intergenerational plasticity and/or genetic effects. For the offspring, however, we conducted a reciprocal common garden experiment mimicking natural winter conditions of the core (warm) and edge (cold) environments and investigated traits based on their population of origin (core or edge) and the winter treatment (warm or cold), which allows us to partly disentangle the contributions of genetics and environment. We are aware that while we tend to interpret origin effects as genetic adaptation, they could represent transgenerational plasticity. Reduced offspring body size and mass and even lower condition for edge origin spiders might represent a fecundity constraint of the mothers, through lower nutrient availability and therefore lower energetic investment into offspring at the edge of the range. Alternatively, the overall size reduction might reveal an adaptation to cold temperatures because the probability of internal ice formation increases with the number of water molecules in a body; thus less body material can improve cold tolerance (Lee, 2010).

The general pattern in our results, for all investigated traits except supercooling strategy, is a strong role of phenotypic plasticity that increases cold tolerance in the cold winter treatment. Overall, the extent of cold tolerance in this species is remarkable, with some core origin spiderlings surviving temperatures as cold as −25°C when cold-acclimated, while edge origin spiderlings can survive temperatures as cold as or colder than −32°C. The presence of this degree of cold tolerance even in the core of the range, where such temperatures do not occur, could suggest that cold tolerance has been maintained in these spiders since the last glacial maximum. At the edge of the range in Estonia, low temperature extremes do occasionally occur, where air temperatures dip below −30°C when Arctic cold fronts move across the country, which could impact the overwintering spiders if they are not covered by snow. Our results show that the colder winters at the edge of the range exert a survival constraint on spiderlings from both origins that survive the cold treatment at lower proportions than the warm treatment with no difference between origins. Recalling that edge spiderlings have lower body condition (**Supplementary Figure S4B**), the similar survival probability compared to the core spiderlings suggests slightly further enhanced cold tolerance in edge origin spiderlings. This assumption is corroborated particularly by their lower supercooling points and lower lethal temperatures. The supercooling point is a highly relevant survival metric for this species since *A. bruennichi* has a freeze avoidant cold tolerance strategy (**Supplementary Text S3**), meaning that they die at the point of internal ice formation. Edge spiderlings consistently have a supercooling point around 1°C lower than core spiderlings (**Figure 3C**). Spiderlings from the cold winter treatment have “low group” supercooling points around 0.5°C lower than spiderlings from the warm winter treatment (**Figure 3C**). These differences are significant, but minimal in effect. However, when considering the ratio of low group to high group supercooling points, e.g. the “supercooling strategy”, the results are more striking: edge spiderlings do not demonstrate plasticity in their supercooling strategy, maintaining low supercooling points regardless of the winter treatment, while core spiderlings in the warm winter treatment have a majority of “high group” supercooling points (**Figure 3D**).

Our analysis of the spiderling metabolome at the end of winter revealed 46 metabolites. We were primarily interested in metabolites which might play a role in enhancing cold tolerance, thus we focused our interpretation on the 10 metabolites that were found in higher quantities in the cold winter treatment and/or edge spiderlings. The accumulation of metabolic intermediates, such as TCA cycle and glycolysis intermediates, aligns with an assumption of a reduced metabolic rate in the cold treatment. It is not known if spiders enter diapause, or if their decreased metabolic rate represents quiescence due only to the effects of low temperature. At the depths of winter, the spiderlings start to move after a short period of warming (around 20 minutes at room temperature; Sheffer, personal observation), suggesting a quiescent state rather than true diapause. Two other groups of metabolites were found in higher relative concentration in the cold treatment and edge spiderlings: amino acids and one cyclic polyol, myo-inositol. Cyclic polyols and free amino acids are known in other arthropods to both colligatively (concentration-dependent) and non-colligatively (concentration independent) increase cold tolerance (e.g. Toxopeus et al., 2019). Cyclic polyols are well known to serve as cryoprotectants, when found in high concentrations. With the relative quantification method we used, we cannot establish whether myo-inositol – already known to occur in high concentrations another spider species (Tanaka, 1995) – is a colligative or non-colligative cryoprotectant, although we expect that it does perform a cryoprotective function.

We confirmed that northeastern populations are genetically differentiated from southwestern populations and discovered that the turnover from genetically “ancestral” genotypes to the “expanding” genotypes occurs in central Germany, as expected based on prior work (Krehenwinkel et al., 2015; Krehenwinkel & Tautz, 2013). We found that the turnover is quite pronounced over a short geographic distance of less than 150 kilometers, despite high dispersal capacity in this species: *A. bruennichi* spiderlings disperse by “ballooning,” a form of aerial dispersal (Follner & Klarenberg, 1995). The genetic differentiation lends support to an argument of genetic adaptation as a contributing factor to the discovered patterns. A detailed investigation into the genes underlying local adaptation requires fuller coverage of the genome than the SNP dataset presented here: our 22,372 SNPs represent only 0.001% of the full genome, with the likelihood of missing coverage of important genes. However, given the results of our exploratory metabolomic analysis, genes related to the synthesis of cryoprotectants such as myo-inositol would be targets of interest.

Further, our findings of several hundred significant genotype-environment associations for environmental variables related to minimum temperature and seasonality lend additional support to the argument that rapid climate-driven genetic adaptation has taken place over the course of this range expansion. In the population genetic and demographic analyses, the three “edge” sites in Estonia emerge as distinct from other northeastern European populations (**Figure 4B, Supplementary Figure S8**), potentially due to observed increases in inbreeding levels at the edge of the range (**Supplementary Figure S7A**). This finding corresponds with the finding by Wolz et al. (2020) of decreased dispersal propensity in edge populations. Decreased dispersal propensity might be a result of selection against leaving the western, coastal part of Estonia either towards the Baltic Sea or further inland, where climatic conditions are much harsher and survival can be assumed unlikely. There and further north, the likelihood of extreme cold snaps nearing and/or exceeding the tolerable temperatures, are high, unless snow cover mitigates the effects of Arctic cold fronts. Since snow cover may not be reliable further north and inland due to climate change, we assume that the range expansion of *A. bruennichi* currently has reached a northern limit.

## Conclusion

We found support for a shared role of phenotypic plasticity and genetic adaptation facilitating establishment of *Argiope bruennichi* spiders in northern climates during the course of a rapid range expansion. Considering adult and juvenile phases of the lifecycle allowed us to detect trait variation that responds to the different selective regimes present in different seasons: in adults at the edge of the range, rapid maturation and smaller adult body size with no concurrent reduction in clutch size or hatching success suggests adaptation to maintain fecundity despite seasonal constraints. In overwintering juveniles, although survival was reduced in the cold winter treatment for both core and edge origins, we found a suite of cold tolerance traits that show both phenotypic plasticity and/or signatures of local adaptation suggesting adaptations to the colder winter temperatures at the northern edge of the distribution after only a few decades of establishment in northern climates. These findings are supported at the genomic level, with a clear differentiation between southern and northern populations and significant genotype-environment associations for environmental variables related to seasonality and minimum winter temperature. Overall, our study sheds light on the potential speed of population changes as more species shift their ranges northward in response to climate change. Further, our findings have implications for how to model organismal responses to climate change: models should include environmental drivers specific to different life stages, allow for both phenotypic plasticity and adaptation in fitness-relevant traits, and model fitness components separately, as seasonality may be a stronger limiting factor for fecundity, while extremes of temperature could exert greater pressure on survival-related traits.

## Data availability

Raw data for the population genomics work is available on NCBI (*Accession Number will be provided upon submission to a peer-reviewed journal*). A file for matching the FASTA file names with the populations as named/numbered in this paper is available as a table in **Supplementary Table S4**. The scripts for filtering SNPs in and outside of exons were written by KJH and can be shared upon reasonable request. Observed means and variances of spider traits have been uploaded to the World Spider Trait Database (*link will be provided upon submission to a peer-reviewed journal*) for utility in meta-analyses, and the raw data can be shared by MMS upon request.

## Acknowledgements

We would like to thank Lydia Smith from the UC Berkeley Evolutionary Genomics Laboratory for her expertise and advice with ddRAD library preparation, as well as the UC Berkeley QB3 Genomics core facility (RRID: SCR_022170) for sequencing services. José Cerca provided helpful troubleshooting advice for bioinformatic processing of the ddRAD data. We are grateful to Cynthia Wang for her work on adapting the library preparation protocol. Many thanks to Heidi Land for her incredible organizational skills helping with the coordination of animal caretaking. We appreciate the dedication of student helpers and interns Alexandra Machnis, Timon Möller, Julia Balk, Anja Junghanns, Kate Miller, and Jasper Murphy. We also thank Marina Wolz for her very helpful advice on the overwintering procedure and how to count and measure offspring, and Barbara Bauer for help with wrangling climate data. Brent Sinclair and Daniel Berner provided valuable suggestions, critique and review of this study as dissertation chapter reviewers. We would like to thank all of the members of the University of Greifswald “Response” graduate school and the General and Systematic Zoology working group for feedback and camaraderie throughout the course of this project for feedback and camaraderie throughout the course of this project. This project was funded by the Deutsche Forschungsgemeinschaft (DFG) as part of the Research Training Group 2010 RESPONSE (GRK 2010) awarded to Gabriele Uhl. Stefan Prost is funded by the University of Oulu and the Research Council of Finland Profi6 336449 programme ‘Biodiverse Anthropocenes’.

## Author Contributions

GU, HK, and MMS designed the study. Samples were collected by GU, HK, LZ, BS, and MMS. DNA was extracted by MMS and HK. MMS performed the library preparation and sequencing, with support and infrastructure from RG. MMS performed genomic analyses with support from SP and KJH. LZ, BS, GU and MMS photographed the spiders in the field. Phenotyping of adult spiders was done by LZ, BS and MMS. The reciprocal common garden experiment and subsequent physiological/phenotypic measurements of offspring were designed and run by MMS, with help from LZ and BS and infrastructure provided by GU and JK. MMS, CA and MW extracted, identified, and analyzed the metabolomic data with input and infrastructure provided by ML. MMS performed statistical analysis of the adult phenotypes, environmental variables, and common garden data, with input from PM and TN. MMS and GU wrote the first drafts of the manuscript. All authors read, commented on, and approved the final draft.

