## Supplementary for "Rapid ecological and evolutionary divergence during a poleward range expansion"

#### Figure S1: Species information for *Argiope bruennichi*

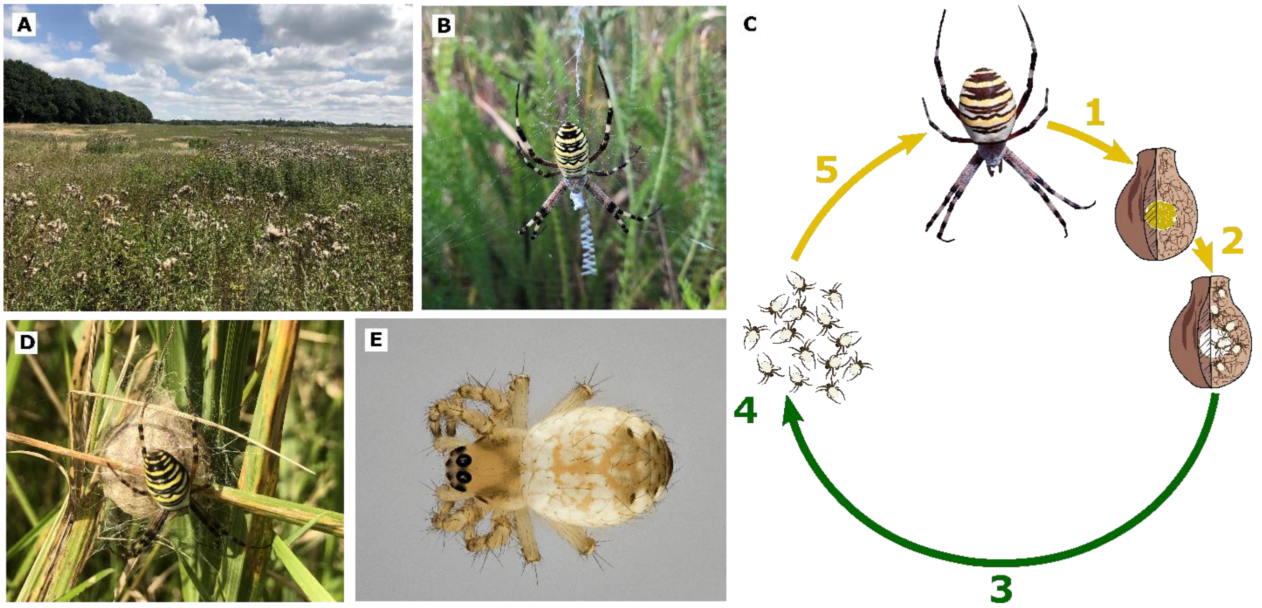

***Argiope bruennichi* species information.** (A) A typical meadow in Germany where *A. bruennichi* individuals are present at high densities. Photo taken in Kemnitz, Germany, by M.M. Sheffer. (B) A female *A. bruennichi* individual in her web, with stabilimentum (zig-zag/zipper-like structure in the web). Photo taken in Kemnitz, Germany, by M.M. Sheffer. (C) *A. bruennichi* life cycle. Spring/Summer months are represented by yellow arrows and numbers, while fall/winter months are represented by green arrows and numbers. 1: Adult females deposit eggs in egg sacs. 2: The eggs hatch and spiderlings molt shortly afterwards. 3: Hatched spiderlings remain in the egg sac over the entirety of winter. 4: In the spring, the spiderlings emerge from the egg sac and disperse. 5: Spiders feed and grow over the course of a few months, maturing and mating toward the end of summer. (D) Adult female with her egg sac, shortly after its construction. Photo taken in Greifswald, Germany, by M.M. Sheffer. (E) Spiderling from southern France removed from an egg sac after winter. Photo taken by T.M. Dederichs.

### Table S1: Sampling sites

**Collecting sites for the phenotypic and genetic aspects studied in the range expansion analysis arranged according to latitude**. The “Sample size” column contains sub-columns for the transect sampling of adult females for genetic analysis (G), adult females for assessing phenotypes (AP), and female of which offspring were used in the common garden experiment (CG). Note that some spiders were used for multiple aspects. Populations with an asterisk indicate populations sampled by Krehenwinkel and colleagues (listed in Krehenwinkel & Tautz 2013).

| Population Number | Population Name | Country | Sample size | | | Latitude (°N) | Longitude (°E) |
|---|---|---|---|---|---|---|---|
| | | | G | AP | CG | | |
| 1 | Belflou | France | 10 | 11 | - | 43.32287 | 1.786019 |
| 2 | Casties | France | 10 | 15 | - | 43.18667 | 1.927365 |
| 3 | Mireval-Lauragais | France | 10 | - | 51 | 43.25554 | 1.963682 |
| 4 | Perry | France | 10 | 10 | 8 | 43.06650 | 2.277690 |
| 5 | Nîmes* | France | 10 | - | - | 43.78000 | 4.400000 |
| 6 | Lyon* | France | 10 | - | - | 45.47968 | 4.830599 |
| 8 | Umkirch | Germany | 10 | 10 | - | 48.05299 | 7.758187 |
| 9 | Kenzingen | Germany | 10 | 7 | - | 48.17536 | 7.783603 |
| 10 | Offenburg | Germany | 10 | 9 | - | 48.44384 | 7.919677 |
| 11 | Brumath | France | 11 | 10 | - | 48.72702 | 7.689943 |
| 12 | Au am Rhein | Germany | 10 | 10 | - | 48.95041 | 8.243427 |
| 13 | Heidelberg | Germany | 10 | 10 | - | 49.42313 | 8.638357 |
| 14 | Gräfenhausen | Germany | 10 | 10 | - | 49.93076 | 8.594512 |
| 15 | Büdingen | Germany | 10 | 10 | - | 50.24868 | 9.078187 |
| 16 | Schotten | Germany | 10 | 4 | - | 50.52840 | 9.117785 |
| 17 | Bad Hersfeld | Germany | 10 | 10 | - | 50.85265 | 9.743685 |
| 18 | Eisenach | Germany | 8 | 3 | - | 51.02786 | 10.35974 |
| 19 | Gierstädt | Germany | 10 | 10 | - | 51.02483 | 10.82997 |
| 20 | Kelbra | Germany | 10 | 10 | - | 51.44596 | 11.10118 |
| 21 | Pegau | Germany | 10 | 10 | - | 51.16905 | 12.25069 |
| 22 | Zeitz | Germany | 4 | 4 | - | 51.97261 | 11.80480 |
| 23 | Barby | Germany | 3 | 3 | - | 51.94881 | 11.91640 |
| 24 | Rabenstein | Germany | 10 | 10 | - | 52.05603 | 12.63610 |
| 25 | Schulzendorf | Germany | 10 | 10 | - | 52.36082 | 13.56633 |
| 26 | Wandlitz | Germany | 10 | 10 | - | 52.74649 | 13.43460 |
| 27 | Templin | Germany | 4 | 4 | - | 53.12951 | 13.48775 |
| 28 | Mittenwald | Germany | 7 | 7 | - | 53.18690 | 13.66299 |
| 29 | Strasburg | Germany | 10 | 11 | - | 53.54765 | 13.72238 |
| 30 | Ludwigsburg | Germany | 10 | 11 | - | 54.11404 | 13.48225 |
| 32 | Grodziszcze* | Poland | 7 | - | - | 52.25839 | 15.56307 |
| 33 | Poznań* | Poland | 10 | - | - | 52.31962 | 17.52766 |
| 34 | Lębork* | Poland | 10 | - | - | 54.54201 | 17.77030 |
| 35 | Palanga* | Lithuania | 10 | - | - | 55.99241 | 21.09721 |
| 36 | Liepāja* | Latvia | 10 | - | - | 56.47952 | 21.02851 |
| 37 | Salaspils* | Latvia | 10 | - | - | 56.85186 | 24.34960 |
| 38 | Ainaži* | Latvia | 10 | - | - | 57.85691 | 24.34681 |
| 39 | Ikla | Estonia | 10 | 10 | 3 | 57.87571 | 24.36059 |
| 40 | Pulgoja | Estonia | 10 | 15 | - | 58.09691 | 24.49626 |
| 41 | Pärnu | Estonia | 10 | 18 | 57 | 58.29375 | 24.59979 |

#### Text S1: Adult phenotypes

##### Dorsal opisthosoma pigmentation of adult females

We used an Olympus Tough F2.0 camera with a ring light flash attachment mounted on a wooden box lined with aluminum foil and blue paper bottom. The spiders were narcotized using $CO_2$ from a soda stream bottle so that we could position them for the photograph in the box. When the camera was mounted, no light could enter the box other than the diffused light from the ring light. The images were processed in Gimp (v.2.10) as follows: we cut out the opisthosoma, added a blue background, and reduced the number of colors in the image to eight, using the "posterize" function. The six colors were then characterized as dark (black, red, green, or violet) or light (white or yellow), and we calculated the number of light and dark pixels per spider in R.

##### Statistical analysis of phenotypes

Body size (tibia-patella length, for adults and offspring) was modeled with a Gaussian distribution, number of eggs laid in an egg sac (clutch size) with a negative binomial distribution with quadratic parameterization ("nbinom2" in R), pigmentation with a betabinomial (light vs dark pixels) distribution, and hatching success of spiderlings from the eggs in an egg sac with a betabinomial distribution. The degree of pigmentation of adult females was modeled with a betabinomial (light vs dark pixels) distribution.

#### Figure S2: Results for additional adult phenotypes

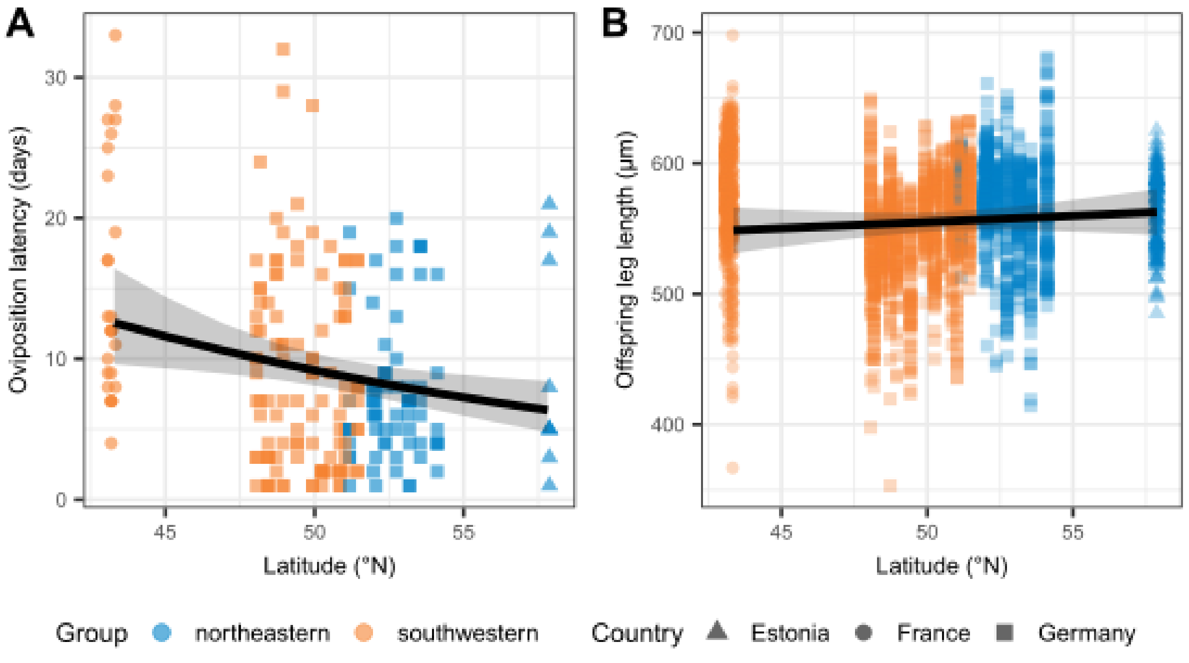

**Figure S2: Phenotypic traits of adult *A. bruennichi* females and their offspring according to latitude.** Points are colored according to their assignment to "southwestern" (orange) or "northeastern" (blue) genetic clusters (see population genetic results, Main text **Figure 4**) and shaped according to the sampling country. Model estimates are plotted as black lines with their 95% confidence intervals as transparent gray ribbons. (**A**) Oviposition latency, the time between collection and oviposition, of adult female spiders, in days. (**B**) Tibia-patella length of the first leg (in µm) of juvenile spiders, a common body size proxy in spiders.

#### Figure S3: Climate simulation of winter regimes applied in the reciprocal common garden experiment

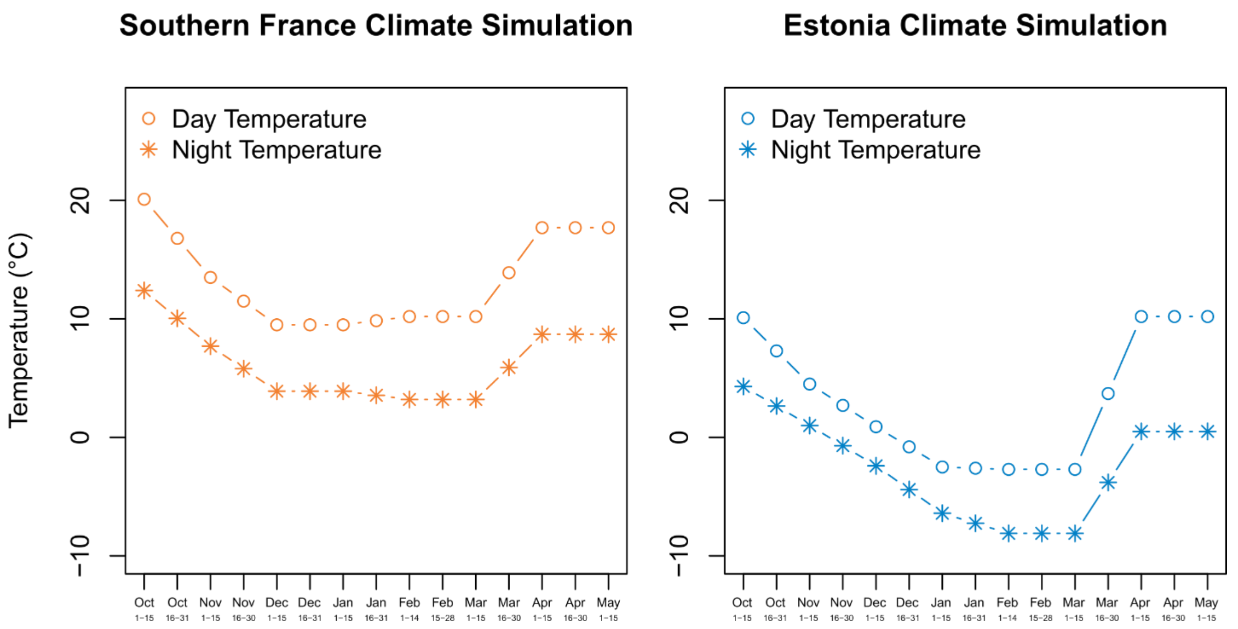

**Figure S2: Climate simulations used for overwintering spiderlings of *A. bruennichi* in the reciprocal common garden experiment.** Egg sacs with spiderlings were placed under either temperature conditions found at the core of the range (Southern France, "warm") or at the edge of the range (Estonia, "cold") (see Figure 1 and Table S1 for sampling sites). Simulations are roughly based on data averaged over 10 years (2004-2014), derived from www.weatheronline.co.uk

---

#### Text S2: Reciprocal common garden experiment

##### Offspring size, mass and condition

After counting the spiderlings in each egg sac, we immediately took the mass of 15 individual spiderlings per clutch using a Sartorius ME5 scale (Sartorius AG, Goettingen, Germany). After weighing, the spiderlings were stored individually in 80% ethanol at room temperature. Later, the first leg on the right side of the spiderling was removed and photographed using a Zeiss Discovery V20 Microscope in combination with a Zeiss AxioCam MRc camera with a 0.63X objective (Zeiss AG, Oberkochen, Germany). From the photos, we measured the length from the proximal end of the patella to the distal end of the tibia ("Tibia-Patella length"), using the Carl Zeiss AxioVision SE64 Software (v.4.9.1.2, Zeiss AG, Oberkochen, Germany). Tibia-patella length is a commonly-used proxy for body size in spiders (Uhl & Vollrath, 2000). To calculate a body condition index, we performed a linear regression of mass against body size, and used the residuals as a response variable, after checking for heteroskedasticity. The pattern of results for body mass and body condition as response variables did not differ, so we include only mass in the main text, but show the graph for leg length and body condition depending on treatment and origin here (Figure S4).

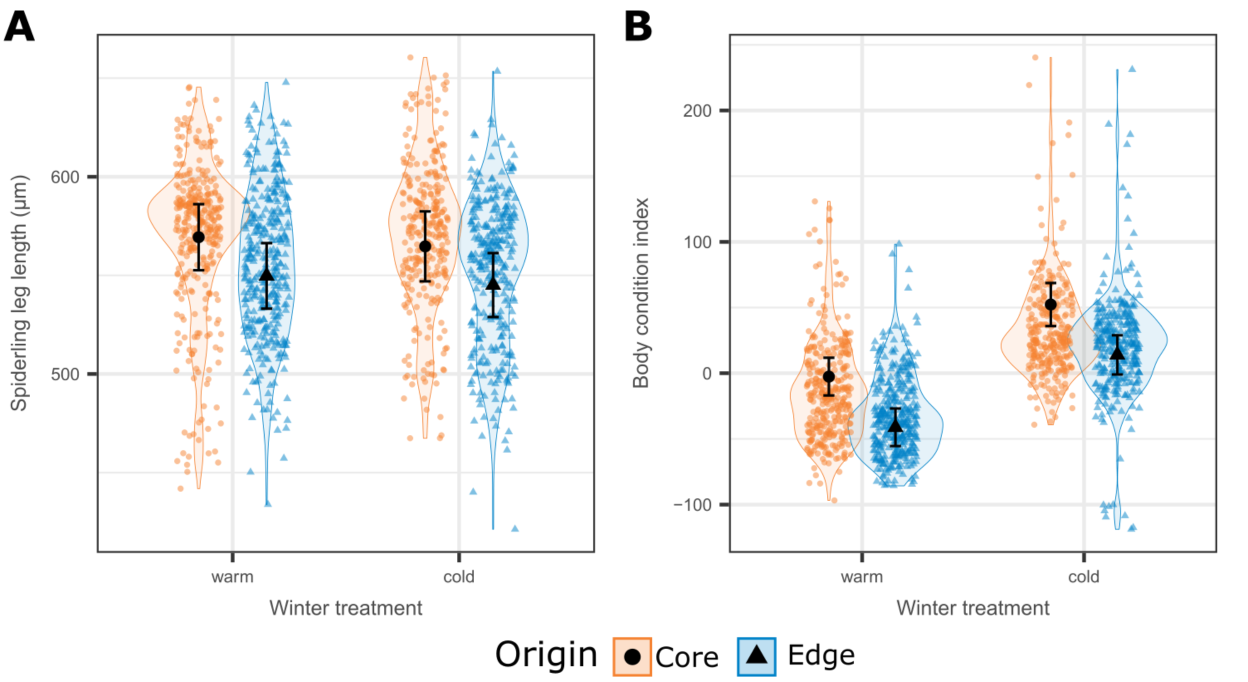

**Figure S4: Body size and condition index** according to winter treatment and origin. Violin plots represent the distribution of the raw data; small points with color according to origin represent the raw data points, jittered to allow visualization of overlapping points. The shape of the points additionally represents the origin. Black points with error bars represent estimated marginal means and 95% confidence intervals for discrete predictor variables. Model predictions are back-transformed to the observed scale. (**A**) Juvenile spider "spiderling" leg length (tibia and patella of the first right leg), a common body size proxy. (**B**) Spiderling body condition, calculated as the residuals of a linear regression of mass and leg length.

#### Cold tolerance experiments after reciprocal common garden

Spiderlings from the four experimental combinations (core/warm, core/cold, edge/warm, edge/cold) were subjected to cold tolerance experiments: lower lethal temperature measurements (20 spiderlings per egg sac), chill coma recovery time (10 spiderlings per egg sac) and supercooling point measurements (8 spiderlings per egg sac). In some families with low survival, small clutch size, or low hatching success, slightly fewer spiderlings were used in some cold tolerance experiments. Details of the methods and protocols are given here.

##### Lower lethal temperature

Of the 20 spiderlings taken from each clutch, 10 were used as test subjects, and 10 were used as a control group, to account for mortality unrelated to the test. The tube containing test spiderlings was placed into a freezer at either -10°C, -18°C, -26°C, or -32°C, for one hour, then placed into a climate chamber held at 15°C, 80% humidity, to recover. The control spiderlings were handled in the same way (carried to the freezer, but not placed inside, and then carried to the climate chamber), but remained at the recovery temperature for the duration of the test (2 hours). After the recovery period, the test and control tubes were opened (blind to treatment, ID, and test/control status) and the number of live and dead spiderlings was recorded. A spiderling was considered dead only if it did not respond to any stimulus (touch with a paintbrush or submersion in ethanol). Of the 1,304 spiderlings in control groups, only 22 died, so we did not incorporate the control data into our analysis.

**Chill coma recovery time**

10 spiderlings per clutch were placed individually into a 3D-printed microwell plate. The wells were painted bright white with non-toxic water-based paint to increase the contrast between the well and the spiderlings. The wells were covered with a glass slide to prevent the spiderlings from escaping, and then the plate was placed into a -9°C freezer for one hour, to induce chill coma. This temperature and exposure time was chosen by preliminary trials, where we found it was sufficient to suppress all movement for some time even after warming, but was not lethal. After one hour, the plate was carefully removed from the freezer, and placed into a climate chamber (15°C, 80% humidity). The glass slide was changed to prevent fogging, and then the spiderlings were filmed in the plate for one hour using an iPhone 6 camera. Analysis of the videos was done manually, with the IDs obscured, by scanning through the video while watching a single spider, until the first movement. Then the video was re-watched at normal speed from a few seconds before the first movement, and the time of the very first movement (i.e. a leg twitch, shift in body position) was marked down as the chill coma recovery time, in seconds. A subset of the videos was re-watched to determine the repeatability of the measurements; the CCR time did not differ significantly between the first and second watching.

**Supercooling point**

The temperature at which the body water freezes can be detected by the heat of crystallization released upon ice formation (Sinclair, Coello Alvarado & Ferguson, 2015). In order to determine the supercooling points of *A. bruennichi* spiderlings depending on winter treatment and origin, eight per clutch were placed individually into a pipette tip, which had been sealed at the tip. A thermocouple (Type K) was inserted into the pipette tip until it came into contact with the spiderling, and was then held in place with a piece of cotton, so that neither the spiderling nor the thermocouple could move. The pipette tips were placed into well-fitting holes drilled into a custom-made aluminum block, and the block was then placed into a dynamic climate chamber (Modell MKT 115, Binder GmbH, Tuttlingen, Germany). Temperature data from the thermocouples was gathered by Pico USB TC-08 thermocouple data loggers (Pico Technology, Cambridgeshire, United Kingdom) once every second for the duration of the trial. Trials started at 5°C and ramped down at a rate of approximately -0.4°C/minute for a total of 150 minutes. The supercooling point was determined as the temperature immediately before a temperature increase above the threshold of 0.3°C/second. In preliminary trials, this threshold turned out to be sensitive enough to detect the more gradual peaks, but not so sensitive that slight background changes in temperature were recorded as supercooling points. Each batch of peaks with supercooling points marked was visually inspected before being entered into the final datasheet.

#### Statistical analysis of common garden data

**Explanatory variables**

In order to assess phenotypic plasticity and genetic adaptation in the common garden experiment, the explanatory variables of central interest were "origin" (southern France or Estonia) and "winter treatment" (warm or cold). All starting models included the interaction of these variables. For the lower lethal temperature experiments, exposure temperature (-10°C, -18°C, -26°C, or -32°C) was also included as an explanatory variable that could interact with origin or winter treatment. Additional, potentially biologically relevant, variables were oviposition latency (time from collection to oviposition), mother's body size and clutch size. Continuous explanatory variables were centered and scaled to allow for interpretation of effects on the same scale (Schielzeth, 2010).

**Random effects**

When spiders from multiple sampling sites were used, sampling site was included as a random effect. For response variables measured at the individual spiderling level, mother's ID was included as a nested random effect within site – specified in the model as (1 | Site / Mother ID).

**Modeling approach**

We analyzed all common garden data using (generalized) linear mixed models in R, using the package 'glmmTMB' (Brooks *et al.*, 2017). Details of the models for each response variable follow below.

For the survival model, we used a betabinomial distribution with default link. Additionally, we found zero inflation due to oviposition latency (an excess of zero survival in late-laid egg sacs). Therefore, we included oviposition latency as a zero-inflation formula. Mass and body size were modeled using a Gaussian distribution.

The lower lethal temperature data were modeled with a betabinomial distribution and default link. These data were recorded as the number surviving or dying within a subset of one clutch. The exposure temperature was added as an explanatory variable, along with all other aforementioned explanatory variables.

The distribution of chill coma recovery times was highly skewed, and attempts to find an appropriate distribution family and model to handle the skewness proved unsuccessful. Therefore, we log-transformed the raw data, allowing us to model with a Gaussian distribution.

Due to the characteristic multimodal distribution of supercooling points (e.g. Sinclair *et al.*, 2003), we followed the common practice and split the data into "low group" (LG) and "high group" (HG) supercooling points at -25.5°C, chosen according to visual inspection of a break in the distribution of the raw data (Figure S5). The LG-SCPs had a normal distribution, and were modeled as a single response variable with a Gaussian distribution. The HG-SCPs were much rarer, and did not allow for modeling directly. Thus, we calculated the per-clutch ratio of HG:LG SCPs, to test whether any of our treatment groups had a tendency toward more HG-SCPs. This HG:LG comparison was modeled with a binomial distribution and logit link.

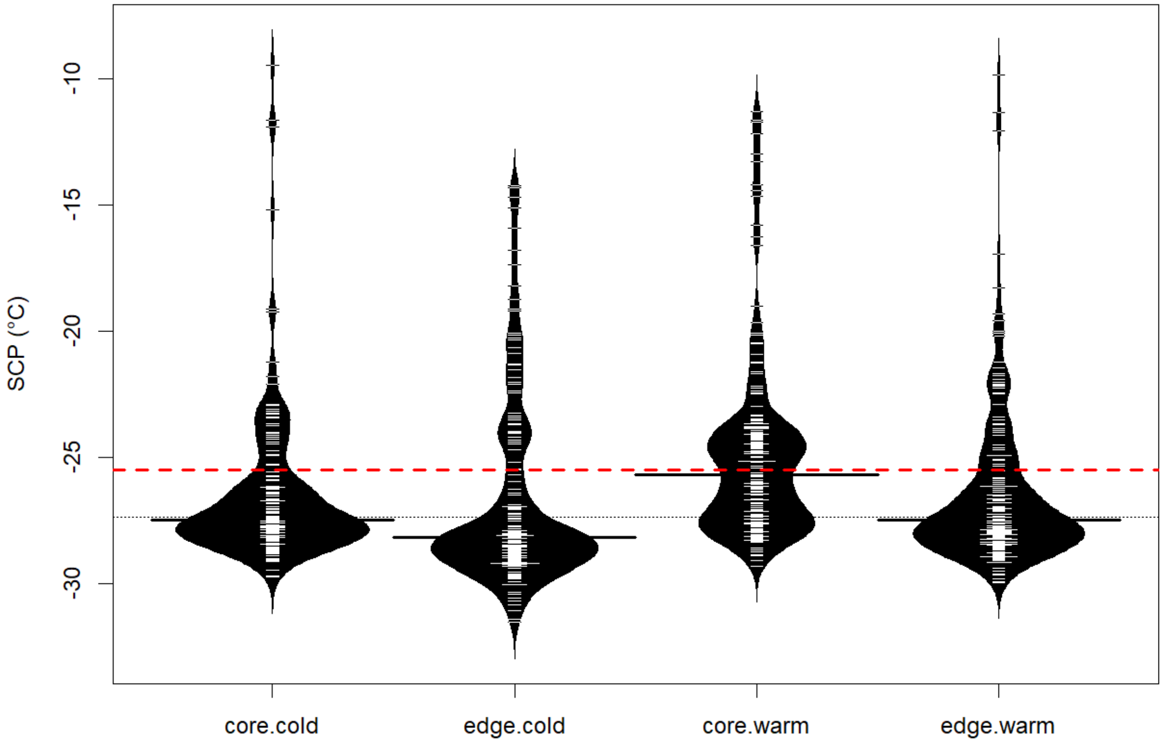

**Figure S5.** "Beanplots" showing the multimodal distribution of supercooling points (SCPs). We split the data into two groups: low group (LG) SCPs, below -25.5°C (dashed red line), and high group (HG) SCPs, above -25.5°C. The LG distribution was Gaussian, and we only used the HG SCPs to calculate a per-family ratio of LG:HG SCPs. White lines represent individual data points, the dotted black line represents the overall median, and the black lines represent the group medians. Groups are labeled on the x-axis according to their origin (core/edge) and winter treatment (cold/warm).

For each full starting model, we performed a backward stepwise model selection using AICc, with random factors always included in the compared models. We did not remove any fixed effects, only the interactions, according to AIC. The model with the lowest AIC was then assessed further. For models with a ΔAIC of less than four, we always proceeded with the model containing maximal interactions within those criteria, as the interaction of origin and winter treatment was of particular interest to us and central to the experimental design. Model assumptions were checked using the R package 'DHARMa', which uses simulated residuals to provide easily interpretable checks of assumptions for many model and distribution types (Hartig, 2021).

Where possible, we used the Nakagawa & Schielzeth method (Nakagawa & Schielzeth, 2013) to calculate a pseudo-$R^2$, implemented in the 'performance' R package (Lüdecke *et al.*, 2021), to assess how much of the variation in the data was explained by the full model (conditional pseudo-$R^2$), and by the model without random effects (marginal pseudo-$R^2$). The Nakagawa & Schielzeth method is not reliable for betabinomial models, thus we used a crude calculation suggested in the 'glmmTMB' FAQs (https://bbolker.github.io/mixedmodels-misc/glmmFAQ.html) to understand the goodness of fit for our betabinomial models (hatching success, overwintering survival, and lower lethal temperature); precisely, we calculated the squared correlation of the predicted values with the observed values of the response variable. This does not provide a comparison of conditional and marginal $R^2$, thus for betabinomial models we only report the conditional $R^2$. We refer to “significant” effects as those where the 95% confidence interval (CI) does not cross zero.

To visualize the results, and present them in the text, we used the ‘ggpredict’ function from the ‘ggeffects’ R package (Lüdecke, 2018) to compute estimated marginal means and 95% CI of the response variable for specific model terms. This allows us to visualize results on the observed scale, making biological interpretation more straightforward. Likewise, although within the model continuous fixed effects were centered and scaled, we plotted the model estimates with the unscaled values.

#### Text S3: Cold tolerance strategy

In order to test if spiderlings survive up to their supercooling point, additional measurements were done in the same way as described above. The only difference was that instead of waiting for a set amount of time, cooling was stopped when approximately half of the spiderlings that were tested at once showed signs of freezing in the live data of the logger (according to Sinclair et al. 2015). Spiderlings were then carefully removed from the pipette tips and placed individually in marked Petri dishes. Survival of the spiderlings was recorded after several minutes at room temperature and merged with the temperature data from the loggers. To test the survival rate of the measurement process itself (e.g. a handling control), a set of experiments were set up, but without ever cooling them: aluminum blocks with spiderlings were kept at room temperature for approximately two hours and survival was assessed in the same way as for the interrupted cooling runs. Experiments were conducted on three separate days with spiderlings from a different egg sac each time.

For the total of 93 tested spiderlings, we categorized the spiderlings either as “frozen dead” (N=44), “frozen survivor” (N=5), “non-frozen dead” (N=6), or “non-frozen survivor” (N=38) (**Figure S6**). These results demonstrate that freezing is lethal, but *A. bruennichi* spiderlings survive up until that point, ergo they employ a freeze-avoidant cold tolerance strategy. The small number of “non-frozen dead” spiderlings can be explained by handling issues when working with such small animals. While it is harder to explain the small number of “frozen survivors,” it is possible that the thermocouples detected a peak from a neighboring sample or from water on the outside of the pipette tip, although we took efforts to avoid that by drying them thoroughly between experiments.

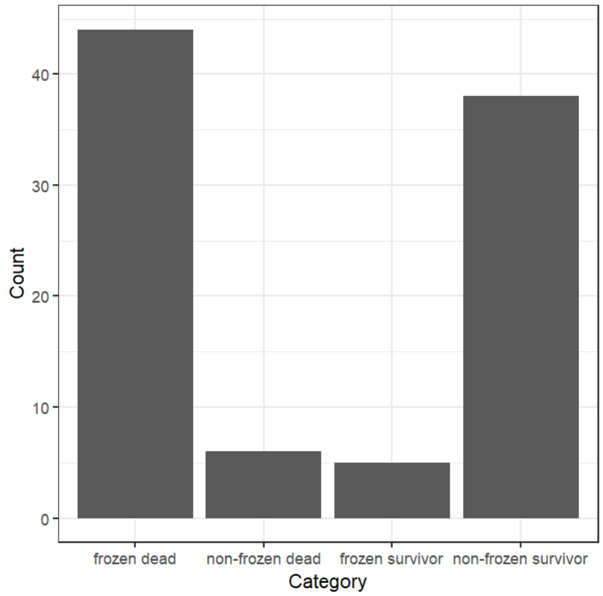

**Figure S6: Cold tolerance strategy.** The x-axis shows the categories into which single spiderlings were categorized, based on the presence or absence of a supercooling peak (peak present = “frozen”, peak absent = “non-frozen”) and whether they did (“survivor”) or did not (“dead”) survive after rewarming. Bars represent the number of individuals in each category.

#### Table S2: Metabolomics

**Table S2: Metabolites and model outputs of differences in relative metabolite concentrations between experimental treatments**. The “Compound” column lists the identity of all 46 individual metabolites identified in our samples. The following five columns represent the model estimate (“Estimate”), standard error (“SE”), sum of squares (Sum Sq), F value, and p-value (Pr>F) of the linear model used to analyze the data. The “Sig” column provides a visual representation of whether the fixed effect in the “Variable” column was significant in the model, with * indicating a p-value < 0.05, ** indicating a p-value < 0.01, and *** indicating a p-value < 0.001.

| Compound | Estimate | SE | Sum Sq | F value | Pr(>F) | Sig | Variable |
|---|---|---|---|---|---|---|---|
| Glyoxylate | 0.157 | 0.208 | 0.725 | 5.203 | 0.034 | * | Winter treatment |
| | 0.087 | 0.199 | 0.542 | 3.890 | 0.063 | | Origin |
| | -0.358 | 0.156 | 0.876 | 6.293 | 0.021 | * | Extractor |
| | 0.367 | 0.305 | 0.201 | 1.447 | 0.243 | | Winter treatment : Origin |
| Pyruvate | 0.241 | 0.270 | 0.122 | 0.516 | 0.481 | | Winter treatment |
| | 0.101 | 0.259 | 0.074 | 0.313 | 0.582 | | Origin |
| | -1.467 | 0.203 | 12.068 | 51.275 | 0.000 | *** | Extractor |
| | -0.422 | 0.397 | 0.266 | 1.130 | 0.300 | | Winter treatment : Origin |
| Lactate | 0.320 | 0.531 | 0.104 | 0.114 | 0.739 | | Winter treatment |
| | 0.481 | 0.510 | 1.937 | 2.130 | 0.160 | | Origin |
| | -3.461 | 0.398 | 67.543 | 74.261 | 0.000 | *** | Extractor |
| | -0.864 | 0.780 | 1.116 | 1.227 | 0.281 | | Winter treatment : Origin |
| Alanine | -0.507 | 0.461 | 0.791 | 1.155 | 0.295 | | Winter treatment |
| | -0.451 | 0.442 | 1.342 | 1.959 | 0.177 | | Origin |

| | | | | | | | |
|---|---|---|---|---|---|---|---|
| | -1.470 | 0.346 | 15.004 | 21.904 | 0.000 | *** | Extractor |
| | 1.649 | 0.676 | 4.071 | 5.943 | 0.024 | * | Winter treatment : Origin |
| B-Hydroxybutyrate | -0.197 | 0.350 | 0.046 | 0.116 | 0.737 | | Winter treatment |
| | -0.358 | 0.336 | 0.404 | 1.023 | 0.324 | | Origin |
| | -0.186 | 0.262 | 0.234 | 0.593 | 0.450 | | Extractor |
| | 0.180 | 0.514 | 0.049 | 0.123 | 0.729 | | Winter treatment : Origin |
| Valine | -0.530 | 0.229 | 1.994 | 11.758 | 0.003 | ** | Winter treatment |
| | 0.389 | 0.220 | 0.956 | 5.637 | 0.028 | * | Origin |
| | -0.331 | 0.172 | 0.615 | 3.624 | 0.071 | | Extractor |
| | -0.094 | 0.337 | 0.013 | 0.078 | 0.783 | | Winter treatment : Origin |
| Urea | -0.112 | 0.207 | 0.075 | 0.545 | 0.469 | | Winter treatment |
| | 0.215 | 0.198 | 0.286 | 2.082 | 0.165 | | Origin |
| | 0.178 | 0.155 | 0.179 | 1.298 | 0.268 | | Extractor |
| | 0.052 | 0.303 | 0.004 | 0.029 | 0.867 | | Winter treatment : Origin |
| Leucine | -0.879 | 0.251 | 3.621 | 17.896 | 0.000 | *** | Winter treatment |
| | 0.012 | 0.240 | 0.114 | 0.564 | 0.461 | | Origin |
| | -0.204 | 0.188 | 0.287 | 1.417 | 0.248 | | Extractor |
| | 0.220 | 0.368 | 0.072 | 0.358 | 0.556 | | Winter treatment : Origin |
| Isoleucine | -0.758 | 0.218 | 2.208 | 14.357 | 0.001 | ** | Winter treatment |
| | -0.056 | 0.210 | 0.068 | 0.445 | 0.513 | | Origin |
| | -0.153 | 0.164 | 0.186 | 1.209 | 0.285 | | Extractor |
| | 0.322 | 0.321 | 0.155 | 1.011 | 0.327 | | Winter treatment : Origin |
| Proline | -0.356 | 0.157 | 0.916 | 11.584 | 0.003 | ** | Winter treatment |
| | 0.328 | 0.150 | 0.670 | 8.474 | 0.009 | ** | Origin |
| | -0.219 | 0.118 | 0.268 | 3.387 | 0.081 | | Extractor |
| | -0.065 | 0.230 | 0.006 | 0.079 | 0.782 | | Winter treatment : Origin |
| Glycine | -1.428 | 0.432 | 2.741 | 4.559 | 0.045 | * | Winter treatment |
| | -0.682 | 0.414 | 0.010 | 0.016 | 0.899 | | Origin |
| | -0.275 | 0.324 | 0.926 | 1.541 | 0.229 | | Extractor |
| | 1.566 | 0.634 | 3.671 | 6.106 | 0.023 | * | Winter treatment : Origin |
| Succinate | 0.808 | 0.262 | 2.436 | 11.059 | 0.003 | ** | Winter treatment |
| | 0.195 | 0.251 | 0.126 | 0.572 | 0.458 | | Origin |
| | -1.607 | 0.196 | 14.263 | 64.742 | 0.000 | *** | Extractor |
| | -0.604 | 0.384 | 0.545 | 2.475 | 0.131 | | Winter treatment : Origin |
| Itaconate | 0.272 | 0.235 | 0.145 | 0.817 | 0.377 | | Winter treatment |
| | 0.710 | 0.225 | 0.718 | 4.047 | 0.058 | | Origin |
| | -0.126 | 0.176 | 0.019 | 0.107 | 0.747 | | Extractor |
| | -0.884 | 0.344 | 1.169 | 6.593 | 0.018 | * | Winter treatment : Origin |
| Fumarate | -1.048 | 0.769 | 9.325 | 4.896 | 0.039 | * | Winter treatment |
| | -0.870 | 0.738 | 7.492 | 3.934 | 0.061 | | Origin |
| | 0.184 | 0.577 | 0.284 | 0.149 | 0.704 | | Extractor |
| | -0.462 | 1.128 | 0.320 | 0.168 | 0.686 | | Winter treatment : Origin |
| Serine | -0.719 | 0.232 | 1.347 | 7.762 | 0.011 | * | Winter treatment |
| | -0.104 | 0.223 | 0.143 | 0.826 | 0.374 | | Origin |
| | -0.264 | 0.174 | 0.540 | 3.109 | 0.093 | | Extractor |
| | 0.505 | 0.341 | 0.382 | 2.201 | 0.153 | | Winter treatment : Origin |

| | | | | | | | |
|---|---|---|---|---|---|---|---|
| Threonine | -0.482 | 0.148 | 1.487 | 21.134 | 0.000 | *** | Winter treatment |
| | 0.294 | 0.142 | 0.608 | 8.646 | 0.008 | ** | Origin |
| | -0.211 | 0.111 | 0.257 | 3.655 | 0.070 | | Extractor |
| | -0.021 | 0.217 | 0.001 | 0.010 | 0.923 | | Winter treatment : Origin |
| B-Alanine | -0.611 | 0.178 | 1.050 | 10.297 | 0.004 | ** | Winter treatment |
| | -0.052 | 0.171 | 0.134 | 1.318 | 0.265 | | Origin |
| | -0.147 | 0.133 | 0.187 | 1.835 | 0.191 | | Extractor |
| | 0.410 | 0.261 | 0.251 | 2.464 | 0.132 | | Winter treatment : Origin |
| Malate | -0.826 | 0.329 | 1.180 | 3.389 | 0.081 | | Winter treatment |
| | -0.413 | 0.315 | 0.002 | 0.006 | 0.938 | | Origin |
| | -1.095 | 0.247 | 7.711 | 22.142 | 0.000 | *** | Extractor |
| | 0.657 | 0.482 | 0.647 | 1.856 | 0.188 | | Winter treatment : Origin |
| Methionine | -0.193 | 0.149 | 0.241 | 3.361 | 0.082 | | Winter treatment |
| | 0.071 | 0.143 | 0.061 | 0.844 | 0.369 | | Origin |
| | -0.421 | 0.112 | 1.016 | 14.175 | 0.001 | ** | Extractor |
| | -0.063 | 0.219 | 0.006 | 0.082 | 0.778 | | Winter treatment : Origin |
| Aspartate | -1.180 | 0.491 | 6.681 | 8.614 | 0.008 | ** | Winter treatment |
| | 0.018 | 0.471 | 0.052 | 0.067 | 0.799 | | Origin |
| | 0.888 | 0.368 | 4.274 | 5.511 | 0.029 | * | Extractor |
| | 0.434 | 0.720 | 0.282 | 0.363 | 0.554 | | Winter treatment : Origin |
| 4-Hydroxyproline | -0.056 | 0.350 | 0.196 | 0.497 | 0.489 | | Winter treatment |
| | 0.359 | 0.336 | 0.407 | 1.031 | 0.322 | | Origin |
| | 0.008 | 0.262 | 0.004 | 0.010 | 0.920 | | Extractor |
| | -0.233 | 0.514 | 0.081 | 0.205 | 0.656 | | Winter treatment : Origin |
| Cysteine | -1.179 | 0.546 | 2.877 | 2.992 | 0.099 | | Winter treatment |
| | -0.120 | 0.524 | 0.886 | 0.922 | 0.348 | | Origin |
| | -0.354 | 0.410 | 1.109 | 1.153 | 0.296 | | Extractor |
| | 1.029 | 0.801 | 1.584 | 1.647 | 0.214 | | Winter treatment : Origin |
| 2-Oxoglutarate | -0.895 | 0.579 | 2.444 | 2.261 | 0.148 | | Winter treatment |
| | 0.159 | 0.556 | 2.415 | 2.234 | 0.151 | | Origin |
| | -2.920 | 0.434 | 50.523 | 46.738 | 0.000 | *** | Extractor |
| | 0.183 | 0.850 | 0.050 | 0.046 | 0.832 | | Winter treatment : Origin |
| Phenylpyruvate | 0.163 | 0.128 | 0.189 | 3.606 | 0.072 | | Winter treatment |
| | -0.171 | 0.122 | 0.134 | 2.549 | 0.126 | | Origin |
| | -0.302 | 0.096 | 0.525 | 10.025 | 0.005 | ** | Extractor |
| | -0.034 | 0.187 | 0.002 | 0.033 | 0.858 | | Winter treatment : Origin |
| Phosphoenolpyruvate | -0.450 | 0.414 | 0.000 | 0.001 | 0.977 | | Winter treatment |
| | -0.343 | 0.397 | 0.056 | 0.102 | 0.753 | | Origin |
| | -0.409 | 0.310 | 1.338 | 2.427 | 0.135 | | Extractor |
| | 0.878 | 0.607 | 1.153 | 2.090 | 0.164 | | Winter treatment : Origin |
| Glutamate | -0.786 | 0.197 | 1.062 | 8.496 | 0.009 | ** | Winter treatment |
| | -0.142 | 0.189 | 0.368 | 2.944 | 0.102 | | Origin |
| | -0.503 | 0.148 | 1.844 | 14.750 | 0.001 | ** | Extractor |
| | 0.729 | 0.289 | 0.795 | 6.361 | 0.020 | * | Winter treatment : Origin |
| Asparagine | -1.616 | 0.614 | 4.147 | 3.418 | 0.079 | | Winter treatment |
| | -0.561 | 0.589 | 0.266 | 0.220 | 0.644 | | Origin |

| | | | | | | | |
|---|---|---|---|---|---|---|---|
| | -0.377 | 0.460 | 1.501 | 1.237 | 0.279 | | Extractor |
| | 1.641 | 0.900 | 4.028 | 3.320 | 0.083 | | Winter treatment : Origin |
| Ribitol | 1.099 | 0.628 | 1.797 | 1.415 | 0.248 | | Winter treatment |
| | 0.201 | 0.602 | 0.685 | 0.539 | 0.471 | | Origin |
| | 0.049 | 0.471 | 0.120 | 0.095 | 0.762 | | Extractor |
| | -1.203 | 0.921 | 2.165 | 1.704 | 0.207 | | Winter treatment : Origin |
| Dihydroxyacetone phosphate | -0.765 | 0.533 | 1.810 | 1.979 | 0.175 | | Winter treatment |
| | 0.442 | 0.511 | 3.302 | 3.611 | 0.072 | | Origin |
| | -0.761 | 0.400 | 3.699 | 4.044 | 0.058 | | Extractor |
| | 0.426 | 0.782 | 0.272 | 0.298 | 0.591 | | Winter treatment : Origin |
| Aconitate | -0.687 | 0.175 | 1.804 | 18.337 | 0.000 | *** | Winter treatment |
| | -0.190 | 0.168 | 0.053 | 0.536 | 0.473 | | Origin |
| | 0.503 | 0.131 | 1.317 | 13.394 | 0.002 | ** | Extractor |
| | 0.374 | 0.256 | 0.209 | 2.127 | 0.160 | | Winter treatment : Origin |
| Glutamine | -1.177 | 1.472 | 0.003 | 0.000 | 0.984 | | Winter treatment |
| | 0.582 | 1.412 | 17.681 | 2.532 | 0.127 | | Origin |
| | 0.529 | 1.104 | 0.598 | 0.086 | 0.773 | | Extractor |
| | 2.677 | 2.160 | 10.728 | 1.536 | 0.230 | | Winter treatment : Origin |
| 2-Phosphoglycerate | -0.087 | 0.170 | 0.084 | 0.901 | 0.354 | | Winter treatment |
| | -0.053 | 0.163 | 0.065 | 0.697 | 0.414 | | Origin |
| | 0.250 | 0.127 | 0.375 | 4.047 | 0.058 | | Extractor |
| | -0.035 | 0.249 | 0.002 | 0.020 | 0.888 | | Winter treatment : Origin |
| 3-Phosphoglycerate | -0.118 | 0.209 | 1.275 | 9.088 | 0.007 | ** | Winter treatment |
| | 0.475 | 0.200 | 0.141 | 1.007 | 0.328 | | Origin |
| | 0.294 | 0.157 | 0.700 | 4.988 | 0.037 | * | Extractor |
| | -0.653 | 0.306 | 0.638 | 4.545 | 0.046 | * | Winter treatment : Origin |
| Ornithine | -0.116 | 0.158 | 0.139 | 1.734 | 0.203 | | Winter treatment |
| | 0.109 | 0.151 | 0.053 | 0.665 | 0.424 | | Origin |
| | -0.180 | 0.118 | 0.175 | 2.183 | 0.155 | | Extractor |
| | -0.091 | 0.231 | 0.012 | 0.154 | 0.699 | | Winter treatment : Origin |
| Citrate | -0.993 | 0.158 | 3.437 | 42.768 | 0.000 | *** | Winter treatment |
| | -0.158 | 0.152 | 0.038 | 0.475 | 0.499 | | Origin |
| | -0.093 | 0.118 | 0.105 | 1.300 | 0.268 | | Extractor |
| | 0.512 | 0.232 | 0.393 | 4.886 | 0.039 | * | Winter treatment : Origin |
| Isocitrate | -0.753 | 0.168 | 0.569 | 6.268 | 0.021 | * | Winter treatment |
| | -0.637 | 0.161 | 0.259 | 2.853 | 0.107 | | Origin |
| | -0.402 | 0.126 | 1.298 | 14.299 | 0.001 | ** | Extractor |
| | 0.868 | 0.246 | 1.127 | 12.414 | 0.002 | ** | Winter treatment : Origin |
| Fructose | 0.223 | 0.218 | 0.120 | 0.788 | 0.385 | | Winter treatment |
| | 0.231 | 0.209 | 0.265 | 1.736 | 0.203 | | Origin |
| | -0.657 | 0.163 | 2.384 | 15.642 | 0.001 | *** | Extractor |
| | -0.254 | 0.319 | 0.096 | 0.631 | 0.436 | | Winter treatment : Origin |
| Glucose | 6.399 | 3.098 | 280.546 | 9.074 | 0.007 | ** | Winter treatment |
| | 2.284 | 2.972 | 79.267 | 2.564 | 0.125 | | Origin |
| | -12.303 | 2.323 | 879.445 | 28.445 | 0.000 | *** | Extractor |
| | -0.760 | 4.545 | 0.864 | 0.028 | 0.869 | | Winter treatment : Origin |

| | | | | | | | |
|---|---|---|---|---|---|---|---|
| Lysine | -0.621 | 0.163 | 1.357 | 15.911 | 0.001 | *** | Winter treatment |
| | -0.054 | 0.156 | 0.076 | 0.894 | 0.356 | | Origin |
| | -0.300 | 0.122 | 0.608 | 7.130 | 0.015 | * | Extractor |
| | 0.286 | 0.239 | 0.122 | 1.436 | 0.245 | | Winter treatment : Origin |
| Glucuronate | 0.007 | 0.425 | 0.863 | 1.483 | 0.237 | | Winter treatment |
| | 0.542 | 0.408 | 5.894 | 10.133 | 0.005 | ** | Origin |
| | -0.765 | 0.319 | 3.985 | 6.852 | 0.016 | * | Extractor |
| | 0.753 | 0.623 | 0.849 | 1.460 | 0.241 | | Winter treatment : Origin |
| Tyrosine | 0.219 | 0.245 | 0.142 | 0.732 | 0.402 | | Winter treatment |
| | 0.076 | 0.235 | 0.063 | 0.326 | 0.574 | | Origin |
| | -1.261 | 0.184 | 8.955 | 46.154 | 0.000 | *** | Extractor |
| | -0.321 | 0.360 | 0.154 | 0.795 | 0.383 | | Winter treatment : Origin |
| myo-Inositol | -0.489 | 0.101 | 1.840 | 56.319 | 0.000 | *** | Winter treatment |
| | 0.316 | 0.097 | 0.556 | 17.008 | 0.001 | *** | Origin |
| | -0.348 | 0.076 | 0.664 | 20.312 | 0.000 | *** | Extractor |
| | -0.144 | 0.148 | 0.031 | 0.951 | 0.341 | | Winter treatment : Origin |
| Tryptophan | -1.319 | 0.443 | 7.193 | 11.397 | 0.003 | ** | Winter treatment |
| | -0.056 | 0.425 | 0.350 | 0.555 | 0.465 | | Origin |
| | -0.954 | 0.332 | 5.675 | 8.993 | 0.007 | ** | Extractor |
| | 0.383 | 0.649 | 0.220 | 0.348 | 0.562 | | Winter treatment : Origin |
| Cystine | -1.338 | 1.493 | 8.647 | 1.204 | 0.285 | | Winter treatment |
| | -0.810 | 1.432 | 19.016 | 2.649 | 0.119 | | Origin |
| | -2.354 | 1.119 | 44.568 | 6.208 | 0.022 | * | Extractor |
| | 5.132 | 2.190 | 39.427 | 5.492 | 0.030 | * | Winter treatment : Origin |
| Fructose-6-phosphate | 0.047 | 0.157 | 0.290 | 3.656 | 0.070 | | Winter treatment |
| | 0.477 | 0.151 | 0.287 | 3.618 | 0.072 | | Origin |
| | 0.412 | 0.118 | 1.184 | 14.916 | 0.001 | *** | Extractor |
| | -0.474 | 0.230 | 0.337 | 4.244 | 0.053 | | Winter treatment : Origin |
| Glucose-6-phosphate | 0.089 | 0.190 | 0.335 | 2.878 | 0.105 | | Winter treatment |
| | 0.595 | 0.182 | 0.493 | 4.239 | 0.053 | | Origin |
| | 0.426 | 0.142 | 1.307 | 11.249 | 0.003 | ** | Extractor |
| | -0.589 | 0.279 | 0.520 | 4.470 | 0.047 | * | Winter treatment : Origin |

#### Text S4: Across latitude: population genomics.

We extracted genomic DNA from the four left legs of the adult female *A. bruennichi* spiders using a plate extraction protocol, as follows: the legs were placed into single well of a 96-well plate containing 3mm steel beads, cell lysis buffer (made of 10mM Tris pH 8, 100mM NaCl, 10mM EDTA pH 8, 0.5% SDS, and double distilled water) and Proteinase K. The plate was then shaken in a GenoGrinder 2010 (OPS Diagnostics) at 1,200 hz for 2 minutes for physical tissue lysis, followed by overnight incubation at 55°C. The next day, RNAse A was added to digest RNA. We then added NaCl to precipitate the proteins, and spun the plate down. We retained the supernatant, containing DNA, and discarded the insoluble protein phase. We used isopropanol to precipitate the DNA from the supernatant, then washed the DNA pellet twice with 70% ethanol. The ethanol was allowed to evaporate, leaving only DNA. The extracted DNA was rehydrated in 50 µl of TE Buffer. DNA quantification was carried out using a Qubit fluorimeter (Thermo Scientific, Waltham, USA) according to the manufacturer's protocol.

We prepared dual-indexed libraries using a double digest restriction-enzyme-associated DNA sequencing (ddRADseq) library preparation protocol, based on the Peterson method (Peterson *et al.*, 2012), as adapted in Maas *et al.* (2018), with 500ng of DNA as starting material for most samples. Some samples started with lower input, but library preparation succeeded nonetheless. We digested the DNA using the restriction enzymes NlaIII (frequent cutter) and SbfI-HF (rare cutter). These enzymes were chosen based on their GC content, to avoid over-digestion of the AT-rich genome. We tested the digestion with *in silico* digestions of the *A. bruennichi* genome assembly (Sheffer *et al.*, 2021b) using SimRAD (Lepais & Weir, 2014), as well as testing digestions in the lab according to the Peterson protocol. We indexed each individual with unique barcodes, then pooled the individuals into 19 libraries of 24 individuals each. Another round of indexing was performed, to give each library of individuals a unique identifier. We quantified the indexed libraries using RT-qPCR, then pooled all of the libraries together. We used a Sage Science Pippin Prep to size select 475-696 bp fragments (including internal adapters), with a peak at 570 bp. The fragment size range was confirmed with a Bioanalyzer High Sensitivity chip (Agilent). We then sequenced 150 bp single-end reads on three lanes of Illumina HiSeq 4000 at UC Berkeley's Vincent J. Coates Genomics Sequencing Lab.

#### Sequence processing and SNP filtering

We used Trimmomatic (v.0.39) (Bolger, Lohse & Usadel, 2014) to remove adapter sequences. We mapped the reads of each sample onto the genome assembly using BWA-MEM (v.0.7.12-r1039) with default settings (Li, 2013), then sorted and indexed the mapped reads using SAMtools sort and SAMtools index (SAMtools v.1.3.1) (Li *et al.*, 2009).

We generated a VCF file for all samples using SAMtools mpileup (SAMtools v.1.3.1) (Li *et al.*, 2009). We then filtered the SNP data using SNPcleaner (v.2.24) (Fumagalli *et al.*, 2014). SNPs were retained for downstream analyses if they had a minimum coverage of 3x per individual, no more than 40**%** of missing data for individual sites, and a maximum coverage of 240x per individual. The cutoff for maximum coverage is ~10 times higher than the average coverage per individual, and helps to avoid SNP calls that fall in highly repetitive regions of the genome. SNP calling resulted in 22,372 biallelic SNPs. We used a custom pipeline (see Data Availability section in main text) to further filter the SNPs based on their position inside/outside of exons and introns. Intergenic SNPs, which did not fall within introns or exons, were used for population genomic analyses that call for neutral loci. We will refer to these as "neutral SNPs" henceforth. For analyses which included all data, including those SNPs falling within genes, we will refer to "all SNPs." To 13,832 "neutral" SNPs did not fall within exons or introns.

To account for uncertainty in SNP and genotype calls based on allele counts, which might introduce noise or bias into downstream analyses (Johnson & Slatkin, 2008; Lynch, 2008), we used an empirical Bayesian framework, implemented in ANGSD (analysis of next-generation sequencing data; Korneliussen, Albrechtsen & Nielsen, 2014) to calculate genotype likelihoods instead of generating SNP genotype calls whenever possible.

#### Population genetic structure and demography

Because our sampling covered a broad geographic area, it was important to quantify the underlying demographic patterns and population genetic structure. To this end, we used several methods: using neutral SNPs, we performed a PCA on the genotype likelihoods, as implemented in ngsTools "PCAngsd"

(Fumagalli *et al.*, 2013). Also using the neutral SNPs, we calculated the per-individual inbreeding coefficients, as well as the pairwise relatedness of all individuals included in the study, using ngsF (Vieira *et al.*, 2013) and ngsRelate (Korneliussen & Moltke, 2015) (Figure S7A). Using all SNPs, we calculated the pairwise $F_{ST}$ for each sampling site in ANGSD (Fumagalli *et al.*, 2013). We visualized the $F_{ST}$ results using a heatmap, calculated in R (R Core Team, 2017) (Figure S7B), as well as a neighbor-joining tree calculated in the R package 'ape' (Paradis & Schliep, 2019)**,** visualized with FigTree (Rambaut & Drummond, 2018).

We then performed clustering using individual level assignments to genetic clusters with ngsAdmix (Skotte, Korneliussen & Albrechtsen, 2013). Due to the stochastic nature of simulation-based clustering programs like ngsAdmix and the like (i.e. STRUCTURE and ADMIXTURE (Pritchard, Stephens & Donnelly, 2000; Alexander, Novembre & Lange, 2009)), we ran 50 replicate analyses with values of *K* ranging from *K*=2 to *K*=20. The results of all replicates were summarized using the program CLUMPAK, which identifies the most common results across replicates for each value of *K* (Kopelman *et al.*, 2015). Graphs of *K*=2, *K*=3, and *K*=4 are shown in Figure S8. The summarized data was then used to visualize individual admixture proportions in R. We used the results of the PCA and ngsAdmix analysis to identify larger groups/"populations" within our data.

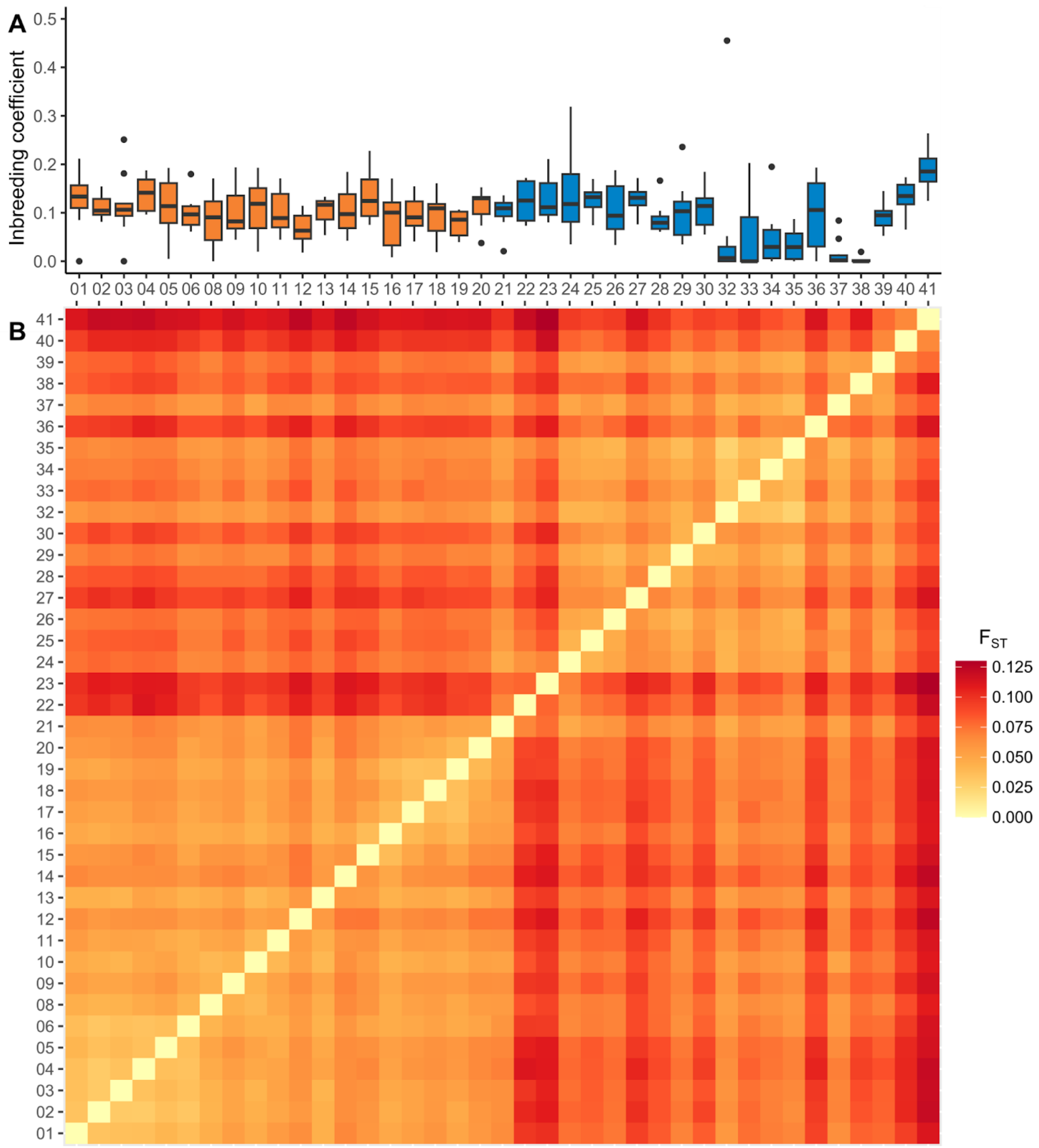

**Figure S7. Demography and differentiation of *A. bruennichi* sampled over a transect from southwestern France to southwestern Estonia (sampling sites: Supplement Table S1).** (A) Boxplots of individual inbreeding coefficients for every sampling site. Boxes encompass the 1st and 3rd quartiles of the data and are colored according to sampling region. Thick horizontal lines represent the median, thin vertical lines represent 1.5 times the upper and lower interquartile range, and black points represent outliers. Box fill color corresponds to the site's assignment to the southwest or northwest genetic cluster. (B) Symmetrical heat-map of pairwise $F_{ST}$ between all sampling sites. The color scale runs from light yellow (lowest differentiation) to dark red (highest differentiation). In both (A) and (B) sites are indicated by numbers, and ordered according to their longitude.

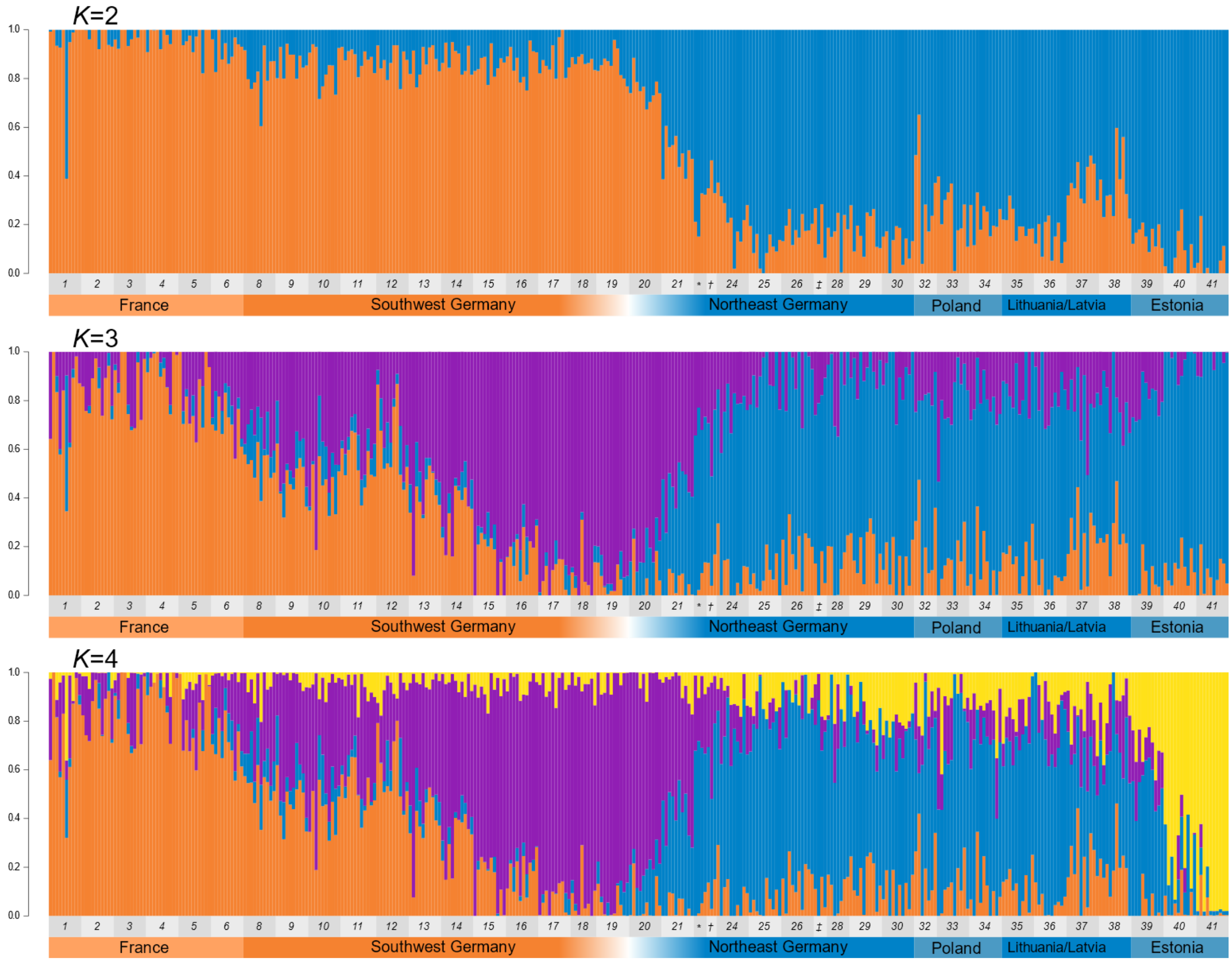

**Figure S8:** Stacked bar plots of *A. bruennichi* individual ancestry assignment, using ngsAdmix results for K=2, K=3, and K=4. Colors represent genetic groupings. Numbers below the bars show the numbered sampling sites from respective countries (Table S1). For sampling sites with few individuals, symbols were used instead of numbers: * represents population number 22, † represents 23, and ‡ represents 27.

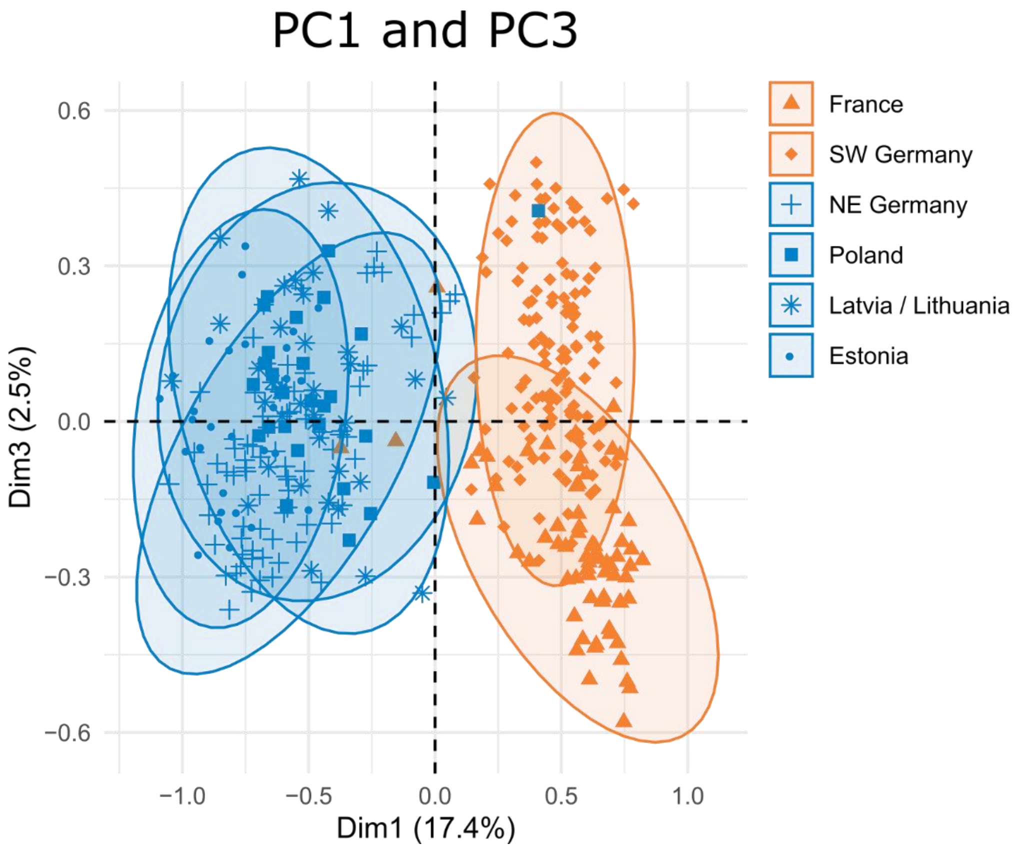

**Figure S9:** Principal component analysis (PCA) using neutral single nucleotide polymorphisms (SNPs), with the first and third principal components plotted. Ellipses represent 95% confidence intervals; points represent individual samples, with shape indicating the sampling country. Points and ellipses are colored according to the site's "southwest" (orange) or "northeast" (blue) assignment.

#### Text S5: Climate data for each site and environmental principal component analysis

We extracted historical climate data for all collecting sites along the transect (Table S1), using an R script and data for the 19 standard bioclimatic variables downloaded from WorldClim at 30 second (~1km$^2$) resolution (Fick & Hijmans, 2017), which include the years 1970-2000. Many of the bioclimatic variables show a distinct change within Germany, although the precise location of the change varies, with northeastern German sites generally having lower temperatures, more seasonality, and less precipitation than southwestern German sites. See Figure S10 for a few select examples of patterns of bioclimatic variables across our sampling sites.

##### Figure S10: Bioclimatic variables across latitude

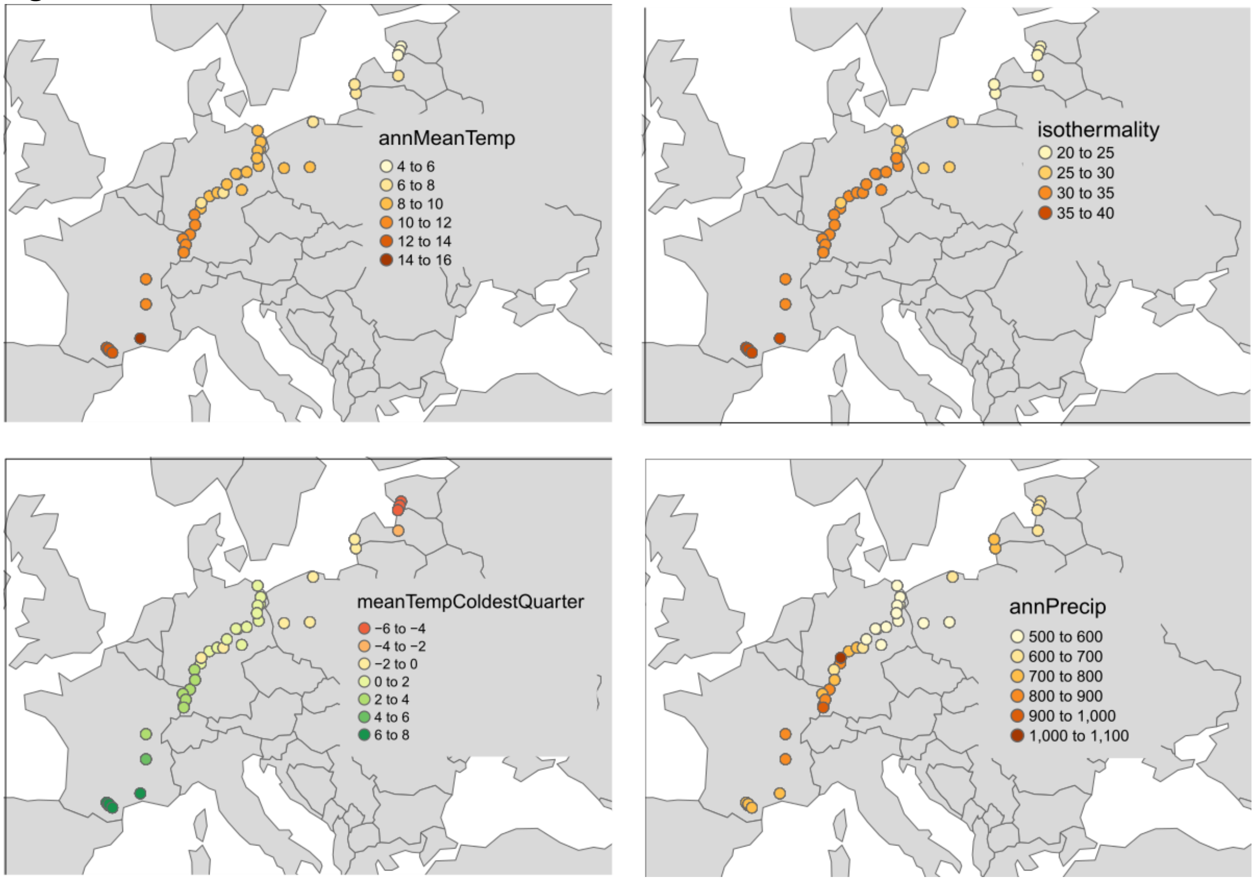

**Figure S10. Select bioclimatic variables at our sampling sites.** Each panel shows a map of Europe, with our sampling sites depicted as points. The bioclimatic variable and its range are overlaid on the graph. annMeanTemp stands for annual mean temperature in °C, meanTempColdestQuarter stands for the mean temperature of the coldest three months of the year in °C, and annPrecip stands for annual precipitation, in centimeters.

To reduce the number of environmental variables in downstream analyses, we summarized the 19 variables into fewer dimensions using a principal component analysis (PCA) using the R package 'FactoExtra' (Kassambara & Mundt, 2020)), following the recommendation of Hoban *et al.* (2016). The first five environmental PCs accounted for 97.6% of the variation in the climate data. PC1 alone accounted for 51.9% of the variation, and varies almost perfectly with latitude (Figure S11). The strongest loadings of this PC were temperature-related, such as the mean temperature of the coldest quarter, the minimum temperature of the coldest month, annual mean temperature, isothermality, and temperature seasonality, although almost all environmental variables had strong contributions to this PC (Figure S12). PC2 accounted for 24.4% of variation, and separated sites more according to precipitation than temperature, with the strongest loadings being the precipitation of the warmest quarter, precipitation of the wettest quarter, annual precipitation, and precipitation of the driest month (Figure S12). PC3 accounted for 11.2% of variation, with the strongest loadings relating to seasonality in temperature and precipitation, as well as the annual temperature range (Figure S13). PC4, which accounted for 6.7% of variation, was strongly related to the mean temperature of the wettest quarter. The strongest loadings for PC5, which accounted for 3.4% of environmental variation, were similar to PC3, but temperature and seasonal precipitation contributed in opposite directions for this PC. See Table S3 for the loadings of each variable for PCs 1-5.

#### Figure S11: Environmental PC1 varies with latitude

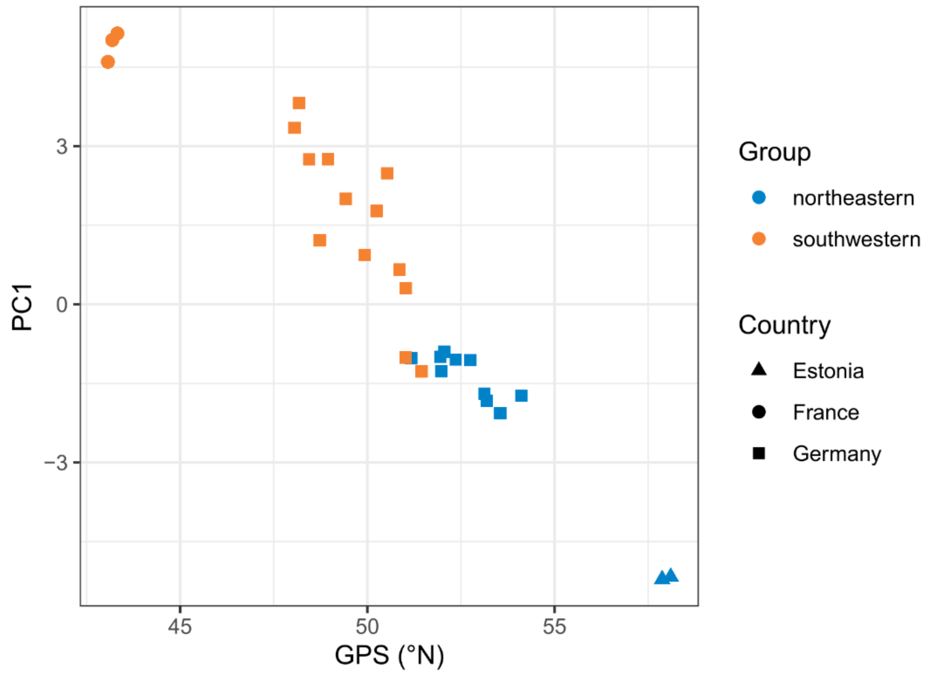

**Figure S11. Environmental PC1 according to latitude.** We summarized 19 “Bioclim” variables into principal components. Points represent single collecting sites, and are colored according to their assignment to “southwestern” (orange) or “northeastern” (blue) genetic clusters. Shapes represent countries.

#### Figure S12: Environmental PCs 1 and 2

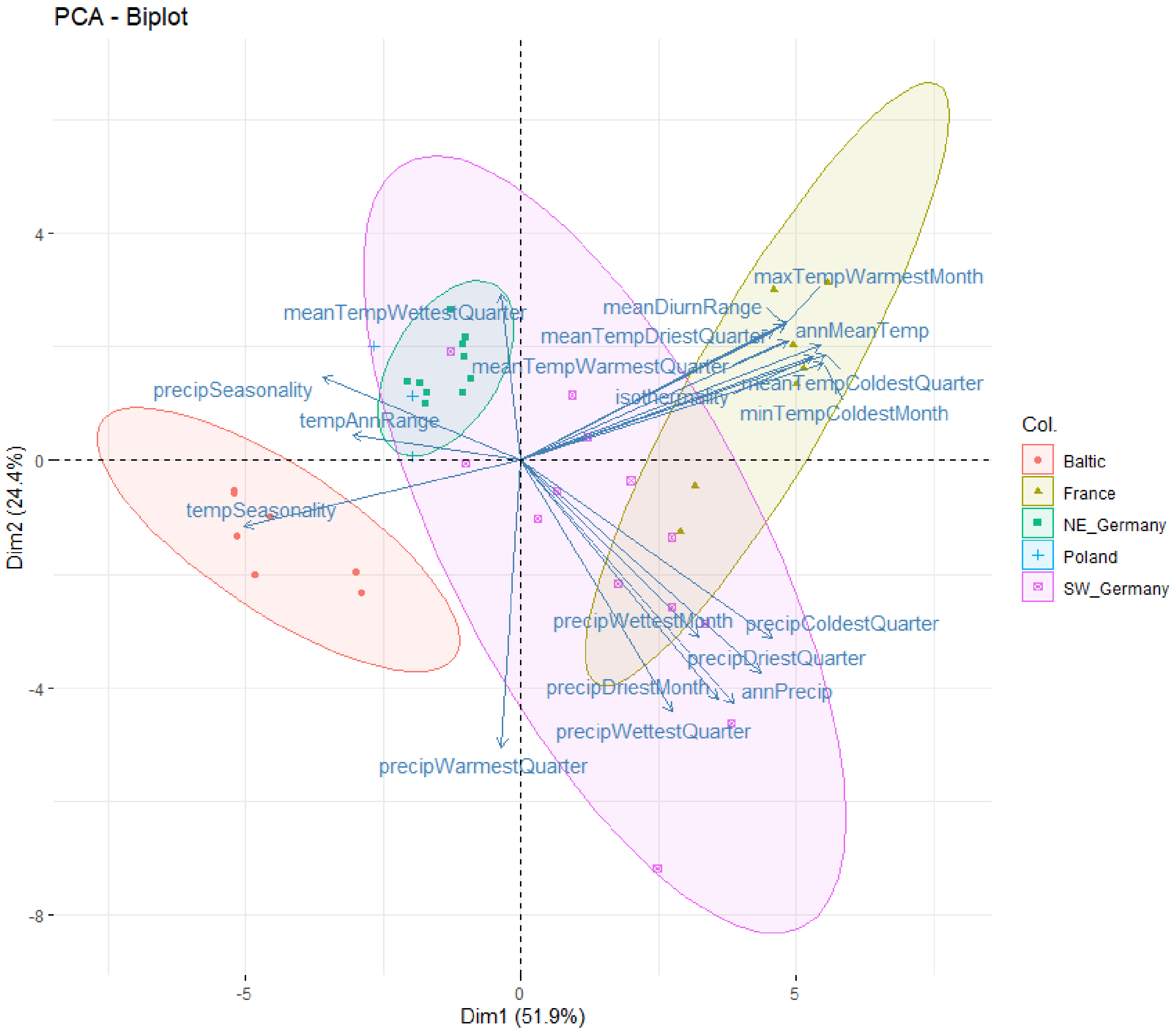

**Figure S12. Biplot of the first two environmental principle components**. We summarized the 19 “Bioclim” variables into principal components, with PCs 1 and 2 visualized here. Point shape and color represents the sampling sub-country, and the ellipses represent the 95% confidence interval of the group. Individual bioclimatic variables and their loadings are depicted as arrows and text overlaying the plot.

#### Figure S13: Environmental PCs 1 and 3

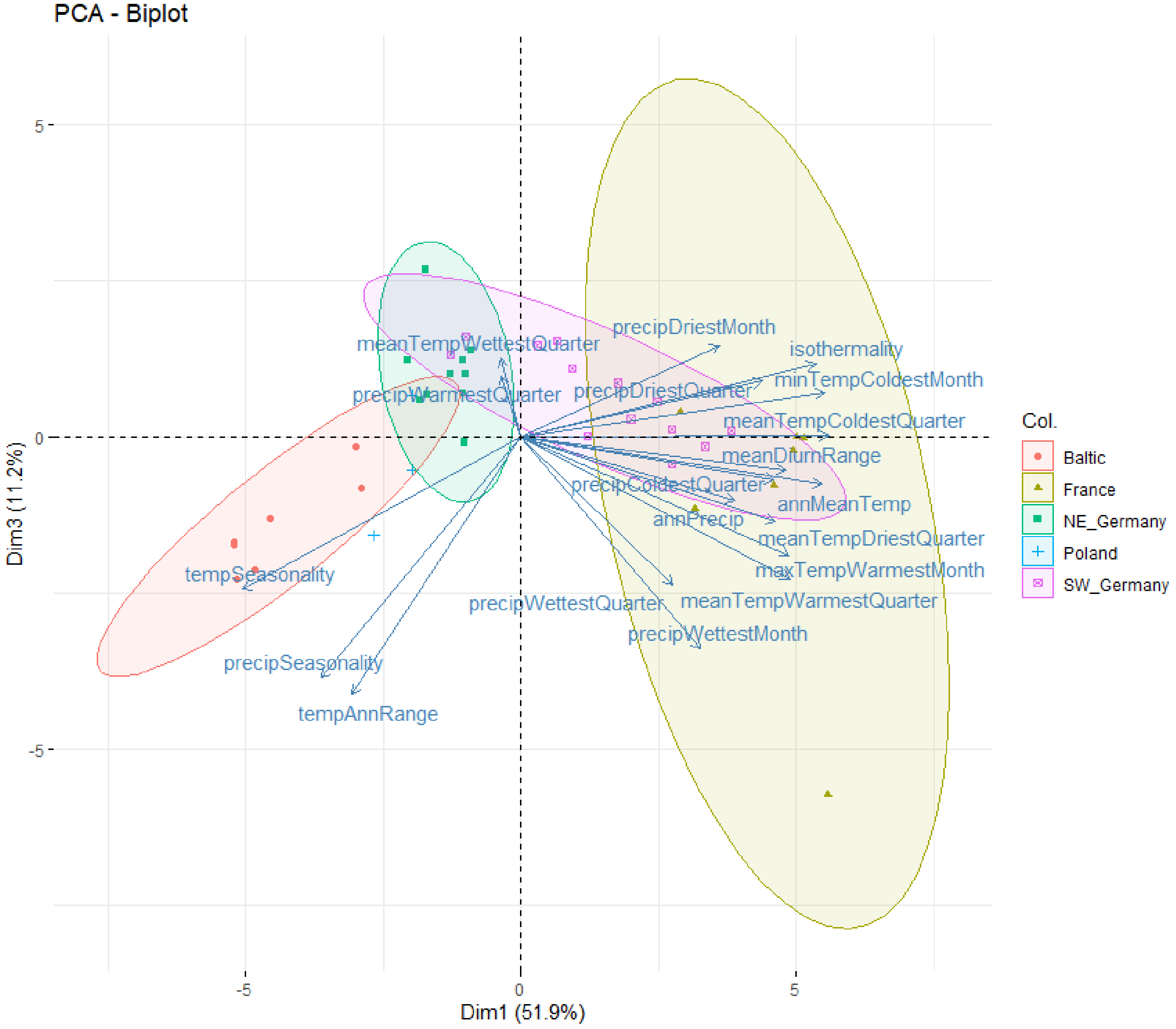

**Figure S13. Biplot of the first and third environmental principle components**. We summarized the 19 “Bioclim” variables into principal components, with PCs 1 and 3 visualized here. Point shape and color represents the sampling sub-country, and the ellipses represent the 95% confidence interval of the group. Individual bioclimatic variables and their loadings are depicted as arrows and text overlaying the plot.

#### Table S3: Loadings of the first five environmental principal components

**Loadings of each bioclimatic variable for the first five principal components**. The first row indicates the percent of the total variation in the data explained by that principal component. The remaining rows represent individual bioclimatic variables. The five strongest contributing variables to each PC are indicated in **bold text**.

| | **PC1** | **PC2** | **PC3** | **PC4** | **PC5** |
|---|---|---|---|---|---|
| **Percent of total variation explained** | 51.9 | 24.4 | 11.2 | 6.7 | 3.4 |
| **annMeanTemp** | **0.293617** | 0.159107 | -0.08674 | -0.04434 | 0.079255 |
| **meanDiurnRange** | 0.258027 | 0.187947 | -0.06097 | **-0.22924** | **-0.3103** |
| **isothermality** | **0.287425** | 0.143882 | 0.136119 | -0.09493 | -0.10269 |
| **tempSeasonality** | **-0.27102** | -0.09048 | -0.281 | -0.13759 | -0.22281 |
| **maxTempWarmestMonth** | 0.261501 | 0.192668 | -0.21764 | -0.16097 | -0.10806 |
| **minTempColdestMonth** | **0.296458** | 0.13427 | 0.083024 | -0.00713 | 0.197998 |
| **tempAnnRange** | -0.1645 | 0.035065 | **-0.47543** | **-0.22438** | **-0.52717** |
| **meanTempWettestQuarter** | -0.0192 | 0.230177 | 0.145362 | **-0.69163** | 0.232633 |
| **meanTempDriestQuarter** | 0.247539 | 0.179178 | -0.15545 | **0.320126** | -0.02145 |
| **meanTempWarmestQuarter** | 0.262044 | 0.166292 | **-0.26135** | -0.11523 | -0.01314 |
| **meanTempColdestQuarter** | **0.299754** | 0.145982 | 0.002638 | 0.031666 | 0.122093 |
| **annPrecip** | 0.208687 | **-0.33588** | -0.11716 | 0.032388 | -0.02936 |
| **precipWettestMonth** | 0.175124 | -0.24308 | **-0.3904** | -0.04958 | **0.28617** |
| **precipDriestMonth** | 0.19303 | **-0.33074** | 0.170342 | -0.17408 | -0.15035 |
| **precipSeasonality** | -0.19419 | 0.116063 | **-0.44472** | 0.022479 | **0.440351** |
| **precipWettestQuarter** | 0.14805 | **-0.34801** | **-0.27309** | -0.09394 | **0.255088** |
| **precipDriestQuarter** | 0.235282 | **-0.29355** | 0.104257 | -0.04437 | -0.1877 |
| **precipWarmestQuarter** | -0.0189 | **-0.3969** | 0.113134 | **-0.3901** | 0.13788 |
| **precipColdestQuarter** | 0.246633 | -0.2459 | -0.07869 | 0.220987 | -0.14219 |

#### Table S4: FASTA file names

A table for matching FASTA file names (base name, e.g. name without file ending, provided in the first column) with information about the sequenced sample. Population number (Pop #) was used in data analysis to order the populations according to their latitude. The Group column provides information on how populations within a country were put together or split into groups for plotting at a coarser level, as in the PCA (Main text Figure 4B, Supplementary Figure S9). Based on genetic analysis, populations were assigned to a still coarser level, as either “northeastern” or “southwestern” genetic clusters, found in the “Assignment to cluster” column.

| FASTA file base name | Pop # | Pop name | Country | Group | Assignment to cluster |
|---|---|---|---|---|---|
| E18-Ik-12-photo | 39 | Ikla | Estonia | Estonia | northeastern |
| E18-Ik-14-photo | 39 | Ikla | Estonia | Estonia | northeastern |
| E18-Ik-18-photo | 39 | Ikla | Estonia | Estonia | northeastern |
| E18-Ik-19-photo | 39 | Ikla | Estonia | Estonia | northeastern |
| E18-Ik-1-photo | 39 | Ikla | Estonia | Estonia | northeastern |
| E18-Ik-2-photo | 39 | Ikla | Estonia | Estonia | northeastern |
| E18-Ik-4-photo | 39 | Ikla | Estonia | Estonia | northeastern |
| E18-Ik-5-photo | 39 | Ikla | Estonia | Estonia | northeastern |
| E18-Ik-7-photo | 39 | Ikla | Estonia | Estonia | northeastern |
| E18-Ik-8-photo | 39 | Ikla | Estonia | Estonia | northeastern |
| E18-Pa-12-photo | 41 | Pärnu | Estonia | Estonia | northeastern |
| E18-Pa-13-photo | 41 | Pärnu | Estonia | Estonia | northeastern |
| E18-Pa-14-photo | 41 | Pärnu | Estonia | Estonia | northeastern |
| E18-Pa-15-photo | 41 | Pärnu | Estonia | Estonia | northeastern |
| E18-Pa-16-photo | 41 | Pärnu | Estonia | Estonia | northeastern |
| E18-Pa-1-photo | 41 | Pärnu | Estonia | Estonia | northeastern |
| E18-Pa-3-photo | 41 | Pärnu | Estonia | Estonia | northeastern |
| E18-Pa-4-photo | 41 | Pärnu | Estonia | Estonia | northeastern |
| E18-Pa-5-photo | 41 | Pärnu | Estonia | Estonia | northeastern |
| E18-Pa-6-photo | 41 | Pärnu | Estonia | Estonia | northeastern |
| E18-PJ-10 | 40 | Pulgoja | Estonia | Estonia | northeastern |
| E18-PJ-13 | 40 | Pulgoja | Estonia | Estonia | northeastern |
| E18-PJ-14 | 40 | Pulgoja | Estonia | Estonia | northeastern |
| E18-PJ-16 | 40 | Pulgoja | Estonia | Estonia | northeastern |
| E18-PJ-18 | 40 | Pulgoja | Estonia | Estonia | northeastern |
| E18-PJ-3 | 40 | Pulgoja | Estonia | Estonia | northeastern |
| E18-PJ-4 | 40 | Pulgoja | Estonia | Estonia | northeastern |
| E18-PJ-6 | 40 | Pulgoja | Estonia | Estonia | northeastern |
| E18-PJ-7 | 40 | Pulgoja | Estonia | Estonia | northeastern |
| E18-PJ-9 | 40 | Pulgoja | Estonia | Estonia | northeastern |
| FrLrLA01 | 6 | Lyon | France | France | southwestern |

| | | | | | |
|---|---|---|---|---|---|
| FrLrLA03 | 6 | Lyon | France | France | southwestern |
| FrLrLA04 | 6 | Lyon | France | France | southwestern |
| FrLrLA05 | 6 | Lyon | France | France | southwestern |
| FrLrLA06 | 6 | Lyon | France | France | southwestern |
| FrLrLA09 | 6 | Lyon | France | France | southwestern |
| FrLrLA10 | 6 | Lyon | France | France | southwestern |
| FrLrLA12 | 6 | Lyon | France | France | southwestern |
| FrLrLA13 | 6 | Lyon | France | France | southwestern |
| FrLrLA15 | 6 | Lyon | France | France | southwestern |
| FrLrNA01 | 5 | Nimes | France | France | southwestern |
| FrLrNA03 | 5 | Nimes | France | France | southwestern |
| FrLrNA04 | 5 | Nimes | France | France | southwestern |
| FrLrNA05 | 5 | Nimes | France | France | southwestern |
| FrLrNA07 | 5 | Nimes | France | France | southwestern |
| FrLrNA08 | 5 | Nimes | France | France | southwestern |
| FrLrNA09 | 5 | Nimes | France | France | southwestern |
| FrLrNA10 | 5 | Nimes | France | France | southwestern |
| FrLrNA11 | 5 | Nimes | France | France | southwestern |
| FrLrNA14 | 5 | Nimes | France | France | southwestern |
| G18-AR-1 | 12 | Au am Rhein | Germany | SW Germany | southwestern |
| G18-AR-10 | 12 | Au am Rhein | Germany | SW Germany | southwestern |
| G18-AR-2 | 12 | Au am Rhein | Germany | SW Germany | southwestern |
| G18-AR-3 | 12 | Au am Rhein | Germany | SW Germany | southwestern |
| G18-AR-4 | 12 | Au am Rhein | Germany | SW Germany | southwestern |
| G18-AR-5 | 12 | Au am Rhein | Germany | SW Germany | southwestern |
| G18-AR-6 | 12 | Au am Rhein | Germany | SW Germany | southwestern |
| G18-AR-7 | 12 | Au am Rhein | Germany | SW Germany | southwestern |
| G18-AR-8 | 12 | Au am Rhein | Germany | SW Germany | southwestern |
| G18-AR-9 | 12 | Au am Rhein | Germany | SW Germany | southwestern |
| G18-BB-1 | 23 | Barby | Germany | NE Germany | northeastern |
| G18-BB-2 | 23 | Barby | Germany | NE Germany | northeastern |
| G18-BB-3 | 23 | Barby | Germany | NE Germany | northeastern |
| G18-BD-11 | 15 | Büdingen | Germany | SW Germany | southwestern |
| G18-BD-12 | 15 | Büdingen | Germany | SW Germany | southwestern |
| G18-BD-14 | 15 | Büdingen | Germany | SW Germany | southwestern |
| G18-BD-15 | 15 | Büdingen | Germany | SW Germany | southwestern |
| G18-BD-19 | 15 | Büdingen | Germany | SW Germany | southwestern |
| G18-BD-3 | 15 | Büdingen | Germany | SW Germany | southwestern |
| G18-BD-4 | 15 | Büdingen | Germany | SW Germany | southwestern |
| G18-BD-5 | 15 | Büdingen | Germany | SW Germany | southwestern |
| G18-BD-7 | 15 | Büdingen | Germany | SW Germany | southwestern |
| G18-BD-9 | 15 | Büdingen | Germany | SW Germany | southwestern |

| | | | | | |
|---|---|---|---|---|---|
| G18-BH-10 | 17 | Bad Hersfeld | Germany | SW Germany | southwestern |
| G18-BH-12 | 17 | Bad Hersfeld | Germany | SW Germany | southwestern |
| G18-BH-17 | 17 | Bad Hersfeld | Germany | SW Germany | southwestern |
| G18-BH-18 | 17 | Bad Hersfeld | Germany | SW Germany | southwestern |
| G18-BH-2 | 17 | Bad Hersfeld | Germany | SW Germany | southwestern |
| G18-BH-3 | 17 | Bad Hersfeld | Germany | SW Germany | southwestern |
| G18-BH-5 | 17 | Bad Hersfeld | Germany | SW Germany | southwestern |
| G18-BH-6 | 17 | Bad Hersfeld | Germany | SW Germany | southwestern |
| G18-BH-8 | 17 | Bad Hersfeld | Germany | SW Germany | southwestern |
| G18-BH-9 | 17 | Bad Hersfeld | Germany | SW Germany | southwestern |
| G18-BM-10 | 11 | Brumath | France | France | southwestern |
| G18-BM-13 | 11 | Brumath | France | France | southwestern |
| G18-BM-15 | 11 | Brumath | France | France | southwestern |
| G18-BM-16 | 11 | Brumath | France | France | southwestern |
| G18-BM-18 | 11 | Brumath | France | France | southwestern |
| G18-BM-19 | 11 | Brumath | France | France | southwestern |
| G18-BM-23 | 11 | Brumath | France | France | southwestern |
| G18-BM-26 | 11 | Brumath | France | France | southwestern |
| G18-BM-3 | 11 | Brumath | France | France | southwestern |
| G18-BM-4 | 11 | Brumath | France | France | southwestern |
| G18-BM-9 | 11 | Brumath | France | France | southwestern |
| G18-BV-1 | 9 | Kenzingen | Germany | SW Germany | southwestern |
| G18-BV-10 | 9 | Kenzingen | Germany | SW Germany | southwestern |
| G18-BV-11 | 9 | Kenzingen | Germany | SW Germany | southwestern |
| G18-BV-12 | 9 | Kenzingen | Germany | SW Germany | southwestern |
| G18-BV-3 | 9 | Kenzingen | Germany | SW Germany | southwestern |
| G18-BV-4 | 9 | Kenzingen | Germany | SW Germany | southwestern |
| G18-BV-5 | 9 | Kenzingen | Germany | SW Germany | southwestern |
| G18-BV-7 | 9 | Kenzingen | Germany | SW Germany | southwestern |
| G18-BV-8 | 9 | Kenzingen | Germany | SW Germany | southwestern |
| G18-BV-9 | 9 | Kenzingen | Germany | SW Germany | southwestern |
| G18-EN-1 | 18 | Eisenach | Germany | SW Germany | southwestern |
| G18-EN-2 | 18 | Eisenach | Germany | SW Germany | southwestern |
| G18-EN-3 | 18 | Eisenach | Germany | SW Germany | southwestern |
| G18-EN-4 | 18 | Eisenach | Germany | SW Germany | southwestern |
| G18-EN-5 | 18 | Eisenach | Germany | SW Germany | southwestern |
| G18-EN-6 | 18 | Eisenach | Germany | SW Germany | southwestern |
| G18-EN-7 | 18 | Eisenach | Germany | SW Germany | southwestern |
| G18-EN-8 | 18 | Eisenach | Germany | SW Germany | southwestern |
| G18-FO-1 | 10 | Offenburg | Germany | SW Germany | southwestern |
| G18-FO-10 | 10 | Offenburg | Germany | SW Germany | southwestern |
| G18-FO-11 | 10 | Offenburg | Germany | SW Germany | southwestern |

| | | | | | |
|---|---|---|---|---|---|
| G18-FO-3 | 10 | Offenburg | Germany | SW Germany | southwestern |
| G18-FO-4 | 10 | Offenburg | Germany | SW Germany | southwestern |
| G18-FO-5 | 10 | Offenburg | Germany | SW Germany | southwestern |
| G18-FO-6 | 10 | Offenburg | Germany | SW Germany | southwestern |
| G18-FO-7 | 10 | Offenburg | Germany | SW Germany | southwestern |
| G18-FO-8 | 10 | Offenburg | Germany | SW Germany | southwestern |
| G18-FO-9 | 10 | Offenburg | Germany | SW Germany | southwestern |
| G18-FP-1 | 21 | Pegau | Germany | NE Germany | northeastern |
| G18-FP-10 | 21 | Pegau | Germany | NE Germany | northeastern |
| G18-FP-11 | 21 | Pegau | Germany | NE Germany | northeastern |
| G18-FP-12 | 21 | Pegau | Germany | NE Germany | northeastern |
| G18-FP-14 | 21 | Pegau | Germany | NE Germany | northeastern |
| G18-FP-15 | 21 | Pegau | Germany | NE Germany | northeastern |
| G18-FP-19 | 21 | Pegau | Germany | NE Germany | northeastern |
| G18-FP-2 | 21 | Pegau | Germany | NE Germany | northeastern |
| G18-FP-4 | 21 | Pegau | Germany | NE Germany | northeastern |
| G18-FP-7 | 21 | Pegau | Germany | NE Germany | northeastern |
| G18-GH-1 | 14 | Gräfenhausen | Germany | SW Germany | southwestern |
| G18-GH-10 | 14 | Gräfenhausen | Germany | SW Germany | southwestern |
| G18-GH-2 | 14 | Gräfenhausen | Germany | SW Germany | southwestern |
| G18-GH-3 | 14 | Gräfenhausen | Germany | SW Germany | southwestern |
| G18-GH-4 | 14 | Gräfenhausen | Germany | SW Germany | southwestern |
| G18-GH-5 | 14 | Gräfenhausen | Germany | SW Germany | southwestern |
| G18-GH-6 | 14 | Gräfenhausen | Germany | SW Germany | southwestern |
| G18-GH-7 | 14 | Gräfenhausen | Germany | SW Germany | southwestern |
| G18-GH-8 | 14 | Gräfenhausen | Germany | SW Germany | southwestern |
| G18-GH-9 | 14 | Gräfenhausen | Germany | SW Germany | southwestern |
| G18-GS-1 | 19 | Gierstädt | Germany | SW Germany | southwestern |
| G18-GS-10 | 19 | Gierstädt | Germany | SW Germany | southwestern |
| G18-GS-11 | 19 | Gierstädt | Germany | SW Germany | southwestern |
| G18-GS-14 | 19 | Gierstädt | Germany | SW Germany | southwestern |
| G18-GS-15 | 19 | Gierstädt | Germany | SW Germany | southwestern |
| G18-GS-16 | 19 | Gierstädt | Germany | SW Germany | southwestern |
| G18-GS-2 | 19 | Gierstädt | Germany | SW Germany | southwestern |
| G18-GS-20 | 19 | Gierstädt | Germany | SW Germany | southwestern |
| G18-GS-5 | 19 | Gierstädt | Germany | SW Germany | southwestern |
| G18-GS-7 | 19 | Gierstädt | Germany | SW Germany | southwestern |
| G18-HB-10 | 13 | Heidelberg | Germany | SW Germany | southwestern |
| G18-HB-12 | 13 | Heidelberg | Germany | SW Germany | southwestern |
| G18-HB-13 | 13 | Heidelberg | Germany | SW Germany | southwestern |
| G18-HB-15 | 13 | Heidelberg | Germany | SW Germany | southwestern |
| G18-HB-17 | 13 | Heidelberg | Germany | SW Germany | southwestern |

| | | | | | |
|---|---|---|---|---|---|
| G18-HB-22 | 13 | Heidelberg | Germany | SW Germany | southwestern |
| G18-HB-3 | 13 | Heidelberg | Germany | SW Germany | southwestern |
| G18-HB-4 | 13 | Heidelberg | Germany | SW Germany | southwestern |
| G18-HB-6 | 13 | Heidelberg | Germany | SW Germany | southwestern |
| G18-HB-7 | 13 | Heidelberg | Germany | SW Germany | southwestern |
| G18-KB-1 | 20 | Kelbra | Germany | NE Germany | southwestern |
| G18-KB-10 | 20 | Kelbra | Germany | NE Germany | northeastern |
| G18-KB-11 | 20 | Kelbra | Germany | NE Germany | northeastern |
| G18-KB-14 | 20 | Kelbra | Germany | NE Germany | northeastern |
| G18-KB-17 | 20 | Kelbra | Germany | NE Germany | northeastern |
| G18-KB-19 | 20 | Kelbra | Germany | NE Germany | northeastern |
| G18-KB-2 | 20 | Kelbra | Germany | NE Germany | northeastern |
| G18-KB-4 | 20 | Kelbra | Germany | NE Germany | northeastern |
| G18-KB-6 | 20 | Kelbra | Germany | NE Germany | northeastern |
| G18-KB-7 | 20 | Kelbra | Germany | NE Germany | northeastern |
| G18-LB-10 | 30 | Ludwigsburg | Germany | NE Germany | northeastern |
| G18-LB-11 | 30 | Ludwigsburg | Germany | NE Germany | northeastern |
| G18-LB-13 | 30 | Ludwigsburg | Germany | NE Germany | northeastern |
| G18-LB-15 | 30 | Ludwigsburg | Germany | NE Germany | northeastern |
| G18-LB-2 | 30 | Ludwigsburg | Germany | NE Germany | northeastern |
| G18-LB-20 | 30 | Ludwigsburg | Germany | NE Germany | northeastern |
| G18-LB-3 | 30 | Ludwigsburg | Germany | NE Germany | northeastern |
| G18-LB-5 | 30 | Ludwigsburg | Germany | NE Germany | northeastern |
| G18-LB-6 | 30 | Ludwigsburg | Germany | NE Germany | northeastern |
| G18-LB-7 | 30 | Ludwigsburg | Germany | NE Germany | northeastern |
| G18-MW-1 | 28 | Mittenwald | Germany | NE Germany | northeastern |
| G18-MW-2 | 28 | Mittenwald | Germany | NE Germany | northeastern |
| G18-MW-3 | 28 | Mittenwald | Germany | NE Germany | northeastern |
| G18-MW-4 | 28 | Mittenwald | Germany | NE Germany | northeastern |
| G18-MW-5 | 28 | Mittenwald | Germany | NE Germany | northeastern |
| G18-MW-7 | 28 | Mittenwald | Germany | NE Germany | northeastern |
| G18-MW-8 | 28 | Mittenwald | Germany | NE Germany | northeastern |
| G18-RS-1 | 24 | Rabenstein | Germany | NE Germany | northeastern |
| G18-RS-11 | 24 | Rabenstein | Germany | NE Germany | northeastern |
| G18-RS-13 | 24 | Rabenstein | Germany | NE Germany | northeastern |
| G18-RS-14 | 24 | Rabenstein | Germany | NE Germany | northeastern |
| G18-RS-16 | 24 | Rabenstein | Germany | NE Germany | northeastern |
| G18-RS-18 | 24 | Rabenstein | Germany | NE Germany | northeastern |
| G18-RS-19 | 24 | Rabenstein | Germany | NE Germany | northeastern |
| G18-RS-3 | 24 | Rabenstein | Germany | NE Germany | northeastern |
| G18-RS-4 | 24 | Rabenstein | Germany | NE Germany | northeastern |
| G18-RS-8 | 24 | Rabenstein | Germany | NE Germany | northeastern |

| | | | | | |
|---|---|---|---|---|---|
| G18-SB-1 | 29 | Strasburg | Germany | NE Germany | northeastern |
| G18-SB-10 | 29 | Strasburg | Germany | NE Germany | northeastern |
| G18-SB-12 | 29 | Strasburg | Germany | NE Germany | northeastern |
| G18-SB-16 | 29 | Strasburg | Germany | NE Germany | northeastern |
| G18-SB-17 | 29 | Strasburg | Germany | NE Germany | northeastern |
| G18-SB-19 | 29 | Strasburg | Germany | NE Germany | northeastern |
| G18-SB-20 | 29 | Strasburg | Germany | NE Germany | northeastern |
| G18-SB-3 | 29 | Strasburg | Germany | NE Germany | northeastern |
| G18-SB-4 | 29 | Strasburg | Germany | NE Germany | northeastern |
| G18-SB-8 | 29 | Strasburg | Germany | NE Germany | northeastern |
| G18-SC-1 | 16 | Schotten | Germany | SW Germany | southwestern |
| G18-SC-10 | 16 | Schotten | Germany | SW Germany | southwestern |
| G18-SC-2 | 16 | Schotten | Germany | SW Germany | southwestern |
| G18-SC-3 | 16 | Schotten | Germany | SW Germany | southwestern |
| G18-SC-4 | 16 | Schotten | Germany | SW Germany | southwestern |
| G18-SC-5 | 16 | Schotten | Germany | SW Germany | southwestern |
| G18-SC-6 | 16 | Schotten | Germany | SW Germany | southwestern |
| G18-SC-7 | 16 | Schotten | Germany | SW Germany | southwestern |
| G18-SC-8 | 16 | Schotten | Germany | SW Germany | southwestern |
| G18-SC-9 | 16 | Schotten | Germany | SW Germany | southwestern |
| G18-SD-1 | 25 | Schulzendorf | Germany | NE Germany | northeastern |
| G18-SD-11 | 25 | Schulzendorf | Germany | NE Germany | northeastern |
| G18-SD-13 | 25 | Schulzendorf | Germany | NE Germany | northeastern |
| G18-SD-15 | 25 | Schulzendorf | Germany | NE Germany | northeastern |
| G18-SD-17 | 25 | Schulzendorf | Germany | NE Germany | northeastern |
| G18-SD-3 | 25 | Schulzendorf | Germany | NE Germany | northeastern |
| G18-SD-5 | 25 | Schulzendorf | Germany | NE Germany | northeastern |
| G18-SD-6 | 25 | Schulzendorf | Germany | NE Germany | northeastern |
| G18-SD-7 | 25 | Schulzendorf | Germany | NE Germany | northeastern |
| G18-SD-8 | 25 | Schulzendorf | Germany | NE Germany | northeastern |
| G18-TL-1 | 27 | Templin | Germany | NE Germany | northeastern |
| G18-TL-2 | 27 | Templin | Germany | NE Germany | northeastern |
| G18-TL-3 | 27 | Templin | Germany | NE Germany | northeastern |
| G18-TL-4 | 27 | Templin | Germany | NE Germany | northeastern |
| G18-UK-11 | 8 | Umkirch | Germany | SW Germany | southwestern |
| G18-UK-15 | 8 | Umkirch | Germany | SW Germany | southwestern |
| G18-UK-17 | 8 | Umkirch | Germany | SW Germany | southwestern |
| G18-UK-18 | 8 | Umkirch | Germany | SW Germany | southwestern |
| G18-UK-2 | 8 | Umkirch | Germany | SW Germany | southwestern |
| G18-UK-3 | 8 | Umkirch | Germany | SW Germany | southwestern |
| G18-UK-5 | 8 | Umkirch | Germany | SW Germany | southwestern |
| G18-UK-6 | 8 | Umkirch | Germany | SW Germany | southwestern |

| | | | | | |
|---|---|---|---|---|---|
| G18-UK-8 | 8 | Umkirch | Germany | SW Germany | southwestern |
| G18-UK-9 | 8 | Umkirch | Germany | SW Germany | southwestern |
| G18-WL-1 | 26 | Wandlitz | Germany | NE Germany | northeastern |
| G18-WL-10 | 26 | Wandlitz | Germany | NE Germany | northeastern |
| G18-WL-12 | 26 | Wandlitz | Germany | NE Germany | northeastern |
| G18-WL-13 | 26 | Wandlitz | Germany | NE Germany | northeastern |
| G18-WL-14 | 26 | Wandlitz | Germany | NE Germany | northeastern |
| G18-WL-2 | 26 | Wandlitz | Germany | NE Germany | northeastern |
| G18-WL-3 | 26 | Wandlitz | Germany | NE Germany | northeastern |
| G18-WL-5 | 26 | Wandlitz | Germany | NE Germany | northeastern |
| G18-WL-7 | 26 | Wandlitz | Germany | NE Germany | northeastern |
| G18-WL-9 | 26 | Wandlitz | Germany | NE Germany | northeastern |
| G18-ZS-1 | 22 | Zeitz | Germany | NE Germany | northeastern |
| G18-ZS-2 | 22 | Zeitz | Germany | NE Germany | northeastern |
| G18-ZS-3 | 22 | Zeitz | Germany | NE Germany | northeastern |
| G18-ZS-4 | 22 | Zeitz | Germany | NE Germany | northeastern |
| LtKlPA01 | 35 | Palanga | Lithuania | Latvia and Lithuania | northeastern |
| LtKlPA02 | 35 | Palanga | Lithuania | Latvia and Lithuania | northeastern |
| LtKlPA03 | 35 | Palanga | Lithuania | Latvia and Lithuania | northeastern |
| LtKlPA04 | 35 | Palanga | Lithuania | Latvia and Lithuania | northeastern |
| LtKlPA05 | 35 | Palanga | Lithuania | Latvia and Lithuania | northeastern |
| LtKlPA06 | 35 | Palanga | Lithuania | Latvia and Lithuania | northeastern |
| LtKlPA07 | 35 | Palanga | Lithuania | Latvia and Lithuania | northeastern |
| LtKlPA08 | 35 | Palanga | Lithuania | Latvia and Lithuania | northeastern |
| LtKlPA09 | 35 | Palanga | Lithuania | Latvia and Lithuania | northeastern |
| LtKlPA10 | 35 | Palanga | Lithuania | Latvia and Lithuania | northeastern |
| LvLiAC01 | 38 | Ainaži | Latvia | Latvia and Lithuania | northeastern |
| LvLiAC02 | 38 | Ainaži | Latvia | Latvia and Lithuania | northeastern |
| LvLiAC03 | 38 | Ainaži | Latvia | Latvia and Lithuania | northeastern |
| LvLiAC04 | 38 | Ainaži | Latvia | Latvia and Lithuania | northeastern |
| LvLiAC05 | 38 | Ainaži | Latvia | Latvia and Lithuania | northeastern |
| LvLiAC06 | 38 | Ainaži | Latvia | Latvia and Lithuania | northeastern |
| LvLiAC07 | 38 | Ainaži | Latvia | Latvia and Lithuania | northeastern |
| LvLiAC08 | 38 | Ainaži | Latvia | Latvia and Lithuania | northeastern |
| LvLiAC12 | 38 | Ainaži | Latvia | Latvia and Lithuania | northeastern |
| LvLiAC13 | 38 | Ainaži | Latvia | Latvia and Lithuania | northeastern |
| LvLpLA01 | 36 | Liepāja | Latvia | Latvia and Lithuania | northeastern |
| LvLpLA02 | 36 | Liepāja | Latvia | Latvia and Lithuania | northeastern |
| LvLpLA03 | 36 | Liepāja | Latvia | Latvia and Lithuania | northeastern |
| LvLpLA08 | 36 | Liepāja | Latvia | Latvia and Lithuania | northeastern |
| LvLpLA10 | 36 | Liepāja | Latvia | Latvia and Lithuania | northeastern |
| LvLpLA12 | 36 | Liepāja | Latvia | Latvia and Lithuania | northeastern |

| | | | | | |
|---|---|---|---|---|---|
| LvLpLA13 | 36 | Liepāja | Latvia | Latvia and Lithuania | northeastern |
| LvLpLA14 | 36 | Liepāja | Latvia | Latvia and Lithuania | northeastern |
| LvLpLA15 | 36 | Liepāja | Latvia | Latvia and Lithuania | northeastern |
| LvLpLA16 | 36 | Liepāja | Latvia | Latvia and Lithuania | northeastern |
| LvSaSA01 | 37 | Salaspils | Latvia | Latvia and Lithuania | northeastern |
| LvSaSA02 | 37 | Salaspils | Latvia | Latvia and Lithuania | northeastern |
| LvSaSA03 | 37 | Salaspils | Latvia | Latvia and Lithuania | northeastern |
| LvSaSA04 | 37 | Salaspils | Latvia | Latvia and Lithuania | northeastern |
| LvSaSA06 | 37 | Salaspils | Latvia | Latvia and Lithuania | northeastern |
| LvSaSA07 | 37 | Salaspils | Latvia | Latvia and Lithuania | northeastern |
| LvSaSA08 | 37 | Salaspils | Latvia | Latvia and Lithuania | northeastern |
| LvSaSA09 | 37 | Salaspils | Latvia | Latvia and Lithuania | northeastern |
| LvSaSA10 | 37 | Salaspils | Latvia | Latvia and Lithuania | northeastern |
| LvSaSA11 | 37 | Salaspils | Latvia | Latvia and Lithuania | northeastern |
| PlGPPA01 | 33 | Poznań | Poland | Poland | northeastern |
| PlGPPA03 | 33 | Poznań | Poland | Poland | northeastern |
| PlGPPA04 | 33 | Poznań | Poland | Poland | northeastern |
| PlGPPA05 | 33 | Poznań | Poland | Poland | northeastern |
| PlGPPA06 | 33 | Poznań | Poland | Poland | northeastern |
| PlGPPA08 | 33 | Poznań | Poland | Poland | northeastern |
| PlGPPA09 | 33 | Poznań | Poland | Poland | northeastern |
| PlGPPA10 | 33 | Poznań | Poland | Poland | northeastern |
| PlGPPA11 | 33 | Poznań | Poland | Poland | northeastern |
| PlGPPA12 | 33 | Poznań | Poland | Poland | northeastern |
| PlLuGA01 | 32 | Grodziszcze | Poland | Poland | northeastern |
| PlLuGA02 | 32 | Grodziszcze | Poland | Poland | northeastern |
| PlLuGA03 | 32 | Grodziszcze | Poland | Poland | northeastern |
| PlLuGA04 | 32 | Grodziszcze | Poland | Poland | northeastern |
| PlLuGA05 | 32 | Grodziszcze | Poland | Poland | northeastern |
| PlLuGA06 | 32 | Grodziszcze | Poland | Poland | northeastern |
| PlLuGA07 | 32 | Grodziszcze | Poland | Poland | northeastern |
| PlPoLA01 | 34 | Lębork | Poland | Poland | northeastern |
| PlPoLA02 | 34 | Lębork | Poland | Poland | northeastern |
| PlPoLA03 | 34 | Lębork | Poland | Poland | northeastern |
| PlPoLA04 | 34 | Lębork | Poland | Poland | northeastern |
| PlPoLA05 | 34 | Lębork | Poland | Poland | northeastern |
| PlPoLA06 | 34 | Lębork | Poland | Poland | northeastern |
| PlPoLA07 | 34 | Lębork | Poland | Poland | northeastern |
| PlPoLA08 | 34 | Lębork | Poland | Poland | northeastern |
| PlPoLA09 | 34 | Lębork | Poland | Poland | northeastern |
| PlPoLA10 | 34 | Lębork | Poland | Poland | northeastern |
| SF18-BN-1 | 1 | Belflou | France | France | southwestern |

| | | | | | |
|---|---|---|---|---|---|
| SF18-BN-12 | 1 | Belflou | France | France | southwestern |
| SF18-BN-14 | 1 | Belflou | France | France | southwestern |
| SF18-BN-18 | 1 | Belflou | France | France | southwestern |
| SF18-BN-3 | 1 | Belflou | France | France | southwestern |
| SF18-BN-4 | 1 | Belflou | France | France | southwestern |
| SF18-BN-5 | 1 | Belflou | France | France | southwestern |
| SF18-BN-7 | 1 | Belflou | France | France | southwestern |
| SF18-BN-8 | 1 | Belflou | France | France | southwestern |
| SF18-BN-9 | 1 | Belflou | France | France | southwestern |
| SF18-CA-10 | 2 | Casties | France | France | southwestern |
| SF18-CA-12 | 2 | Casties | France | France | southwestern |
| SF18-CA-13 | 2 | Casties | France | France | southwestern |
| SF18-CA-14 | 2 | Casties | France | France | southwestern |
| SF18-CA-17 | 2 | Casties | France | France | southwestern |
| SF18-CA-18 | 2 | Casties | France | France | southwestern |
| SF18-CA-4 | 2 | Casties | France | France | southwestern |
| SF18-CA-5 | 2 | Casties | France | France | southwestern |
| SF18-CA-6 | 2 | Casties | France | France | southwestern |
| SF18-CA-7 | 2 | Casties | France | France | southwestern |
| SF18-MiO-12 | 3 | Mireval | France | France | southwestern |
| SF18-MiO-18 | 3 | Mireval | France | France | southwestern |
| SF18-MiO-27 | 3 | Mireval | France | France | southwestern |
| SF18-MiO-40 | 3 | Mireval | France | France | southwestern |
| SF18-MiO-5 | 3 | Mireval | France | France | southwestern |
| SF18-MiW-17 | 3 | Mireval | France | France | southwestern |
| SF18-MiW-23 | 3 | Mireval | France | France | southwestern |
| SF18-MiW-39 | 3 | Mireval | France | France | southwestern |
| SF18-MiW-42 | 3 | Mireval | France | France | southwestern |
| SF18-MiW-8 | 3 | Mireval | France | France | southwestern |
| SF18-PE-12-photo | 4 | Perry | France | France | southwestern |
| SF18-PE-15-photo | 4 | Perry | France | France | southwestern |
| SF18-PE-17-photo | 4 | Perry | France | France | southwestern |
| SF18-PE-1-photo | 4 | Perry | France | France | southwestern |
| SF18-PE-2-photo | 4 | Perry | France | France | southwestern |
| SF18-PE-3-photo | 4 | Perry | France | France | southwestern |
| SF18-PE-4-photo | 4 | Perry | France | France | southwestern |
| SF18-PE-6-photo | 4 | Perry | France | France | southwestern |
| SF18-PE-7-photo | 4 | Perry | France | France | southwestern |
| SF18-PE-8-photo | 4 | Perry | France | France | southwestern |
